# Efficient Bayesian mixed model analysis increases association power in large cohorts

**DOI:** 10.1101/007799

**Authors:** Po-Ru Loh, George Tucker, Brendan K Bulik-Sullivan, Bjarni J Vilhjálmsson, Hilary K Finucane, Daniel I Chasman, Paul M Ridker, Benjamin M Neale, Bonnie Berger, Nick Patterson, Alkes L Price

## Abstract

Linear mixed models are a powerful statistical tool for identifying genetic associations and avoiding confounding. However, existing methods are computationally intractable in large cohorts, and may not optimize power. All existing methods require time cost O(MN^2^) (where N = #samples and M = #SNPs) and implicitly assume an infinitesimal genetic architecture in which effect sizes are normally distributed, which can limit power. Here, we present a far more efficient mixed model association method, BOLT-LMM, which requires only a small number of O(MN) iterations and increases power by modeling more realistic, non-infinitesimal genetic architectures via a Bayesian mixture prior on marker effect sizes. We applied BOLT-LMM to nine quantitative traits in 23,294 samples from the Women’s Genome Health Study (WGHS) and observed significant increases in power, consistent with simulations. Theory and simulations show that the boost in power increases with cohort size, making BOLT-LMM appealing for GWAS in large cohorts.

---

Linear mixed models are emerging as the method of choice for association testing in genomewide association studies (GWAS) because they account for both population stratification and cryptic relatedness and achieve increased statistical power by jointly modeling all genotyped markers [1–12]. However, existing mixed model methods still have limitations. First, mixed model analysis is computationally expensive. Despite a series of recent algorithmic advances, current algorithms require O(MN^2^) total running time (assuming N < M), where M is the number of markers and N is the sample size, a cost that is becoming prohibitive for large cohorts [12]. Second, current mixed model methods fall short of achieving maximal statistical power owing to suboptimal modeling assumptions regarding the genetic architectures underlying phenotypes. The standard linear mixed model implicitly assumes that all variants are causal with small effect sizes drawn from independent Gaussian distributions—the “infinitesimal model”—whereas in reality, complex traits have been estimated to have on the order of only a few thousand causal loci [13, 14].

Methodologically, efforts to more accurately model non-infinitesimal genetic architectures have followed two general thrusts. One approach is to apply the standard infinitesimal mixed model but adapt the input data. For example, large-effect loci can be explicitly identified and conditioned out as fixed effects [7], or the mixed model can be applied to only a selected subset of markers [9, 11, 15, 16]. A more flexible alternative approach is to adapt the mixed model itself by taking a Bayesian perspective and modeling SNP effects with non-Gaussian prior distributions that better accommodate both small- and large-effect loci. Such methods were pioneered in livestock genetics to improve prediction of genetic values [17] and have been extensively developed in the plant and animal breeding literature for the purpose of genomic selection [18]. While modeling methods that improve prediction should in theory enable corresponding improvements in statistical power of association analysis (via conditioning on other associated loci when testing a candidate marker [9, 12]), a challenge of applying Bayesian methods in the GWAS setting is that Bayesian statistics are not readily interpretable in the customary hypothesis testing framework.

Here, we present an algorithm that performs mixed model analysis in a small number of O(MN)-time iterations and increases power by modeling non-infinitesimal genetic architectures. Our algorithm fits a Gaussian mixture model of SNP effects [19], using a fast variational approximation [20–22] to compute approximate phenotypic residuals, and tests the residuals for association with candidate markers via a retrospective score statistic [23] that provides a bridge between Bayesian modeling for phenotype prediction and the frequentist association testing framework. We calibrate our statistic using an approach based on the recently developed LD Score regression technique [24]. The entire procedure operates directly on raw genotypes stored compactly in memory and does not require computing or storing a genetic relationship matrix. In the special case of the infinitesimal model, we achieve results equivalent to existing methods at dramatically reduced time and memory cost.

We provide an efficient software implementation of our algorithm, BOLT-LMM, and demonstrate its computational efficiency on simulated data sets of up to 480,000 individuals. Our simulations also show that BOLT-LMM achieves increased association power over standard infinitesimal mixed model analysis of traits driven by a few thousand causal SNPs. We applied BOLT-LMM to perform mixed model analysis of nine quantitative traits in 23,294 samples from the Women’s Genome Health Study (WGHS) [25] and observed increased association power equivalent to an up to 10% increase in effective sample size. We demonstrate through theory and simulations that the boost in power increases with cohort size, making BOLT-LMM a promising approach for largescale GWAS.

## Results

### Overview of Methods

The BOLT-LMM algorithm consists of four main steps, each of which run in a small number of O(MN)-time iterations. These steps are: (1a) Estimate variance parameters; (1b) Compute infinitesimal mixed model association statistics (denoted BOLT-LMM-inf); (2a) Estimate Gaussian mixture parameters; (2b) Compute Gaussian mixture model association statistics (BOLT-LMM). Step 1a computes results nearly identical to standard variance components analysis but applies a new stochastic approximation algorithm that reduces time and memory cost by circumventing spectral decomposition, which is expensive for large sample sizes. Instead, the approximation algorithm only requires solving linear systems of mixed model equations, which can be accomplished efficiently using conjugate gradient iteration [26, 27]. Step 1b likewise circumvents spectral decomposition by introducing a new retrospective mixed model association statistic similar to GRAMMAR-Gamma [10] and MASTOR [23], which we compute—up to a calibration constant— using only solutions to linear systems of equations. We estimate the calibration constant by computing and comparing the new statistic and the standard prospective mixed model statistic at a random subset of SNPs, which can likewise be accomplished efficiently using conjugate gradient iteration. This procedure is similar in spirit to GRAMMAR-Gamma calibration but requires only O(MN)-time iterations.

Steps 2a and 2b are Gaussian mixture parallels of steps 1a and 1b. BOLT-LMM’s non-infinitesimal model amounts to a relaxation of the standard mixed model, which from a Bayesian perspective imposes a Gaussian prior distribution on SNP effect sizes. BOLT-LMM relaxes this modeling assumption by allowing the prior distribution to be a mixture of two Gaussians, giving the model greater flexibility to accommodate large-effect SNPs while maintaining effective modeling of genome-wide effects (e.g., ancestry). For the Gaussian mixture model, it is no longer tractable to perform exact posterior inference, so BOLT-LMM instead computes a variational approximation [20–22] that converges after a small number of O(MN)-time iterations. Step 2a applies this method within 5-fold cross-validation to estimate best-fit parameters for the prior distribution (taking into account variance parameters estimated in step 1a) based on out-of-sample prediction accuracy. If the prediction accuracy of the best-fit Gaussian mixture model exceeds that of the infinitesimal model by at least a specified amount, step 2b is then run to compute association statistics by testing each SNP against the residual phenotype obtained from the Gaussian mixture model and calibrating the test statistics against the results of step 1b using LD Score regression [24]. Otherwise, the BOLT-LMM association statistic is the same as BOLT-LMM-inf. Both step 1b and step 2b are performed using a leave-one-chromosome-out (LOCO) scheme to avoid proximal contamination [9,12]. Further details of the method are provided in Online Methods and the Supplementary Note. The key properties of BOLT-LMM in terms of speed and model specification are compared to existing mixed model association methods in Table 1.

**Table 1.** Comparison of fast mixed model association methods that model all SNPs.

| Method<sup>a</sup> | Requires $O(MN^2)$ time | Avoids proximal contamination | Models non-infinitesimal genetic architecture |
| --- | --- | --- | --- |
| EMMAX [3] | X |  |  |
| FaST-LMM [9] | X<sup>b</sup> | X |  |
| FaST-LMM-Select [9, 11, 15] | X<sup>b</sup> | X | X<sup>c</sup> |
| GEMMA [6] | X |  |  |
| GRAMMAR-Gamma [10] | X |  |  |
| GCTA-LOCO [12] | X | X |  |
| BOLT-LMM |  | X | X |

<sup>a</sup>For methods that have been updated over multiple publications, we cite and list characteristics of the latest published version. <sup>b</sup>If $M < N$ , FaST-LMM and FaST-LMM-Select can complete in $O(M^2N)$ time. <sup>c</sup>FaST-LMM-Select models non-infinitesimal genetic architectures by restricting the mixed model to a subset of SNPs; a caveat of this approach is that it may incur susceptibility to confounding from stratification [12].

### BOLT-LMM is much more computationally efficient than existing methods

To analyze the computational performance of BOLT-LMM, we simulated data sets of sizes ranging up to N = 3,750 to 480,000 individuals and M = 300,000 SNPs. We used genotypes from the WTCCC2 data set [28] analyzed in ref. [12], which contains 15,633 individuals of European ancestry, to form mosaic chromosomes, and we used a phenotype model in which 5,000 SNPs explained 20% of phenotypic variance (Online Methods).

We benchmarked BOLT-LMM against existing mixed model association methods, running each method for up to 10 days on machines with 96GB of memory. BOLT-LMM completed all analyses through N = 480,000 individuals within these constraits, whereas previous methods could only analyze a maximum of N = 7,500–30,000 individuals (Fig. 1). All previous methods require O(MN^2^) running time, whereas BOLT-LMM requires only *≈*O(MN^1.5^) time (Fig. 1a and Supplementary Fig. 1a). We also observed substantial savings in memory use with BOLT-LMM, which requires little more than the MN/4 bytes of memory needed to store raw genotypes, much less than existing mixed model methods (Fig. 1b and Supplementary Fig. 1b).

**Figure 1.**
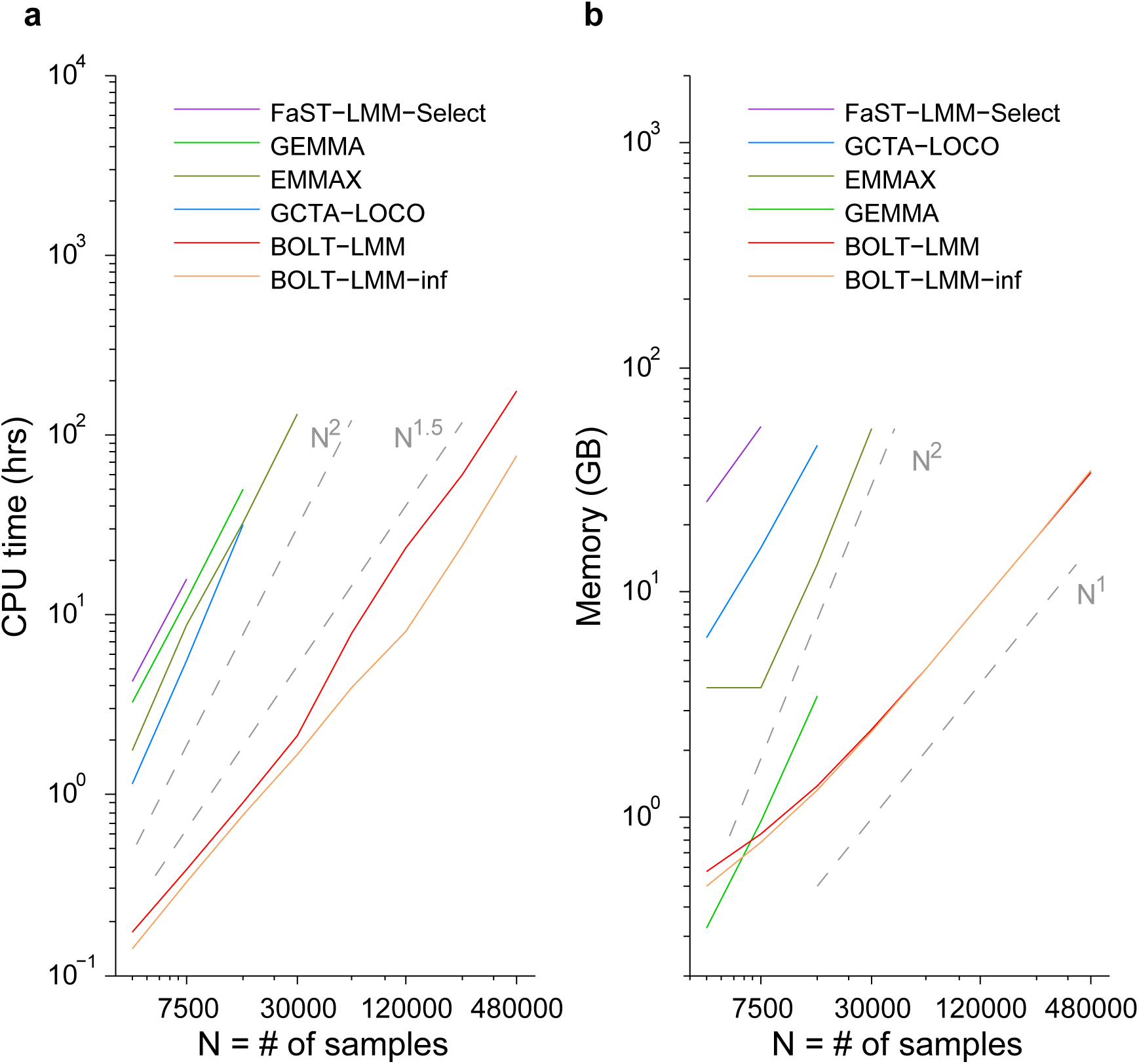
Computational performance of mixed model association methods. Log-log plots of (a) run time and (b) memory as a function of sample size (N). Slopes of the curves correspond to exponents of power-law scaling with N. Benchmarking was performed on simulated data sets in which each sample was generated as a mosaic of genotype data from 2 random “parents” from the WTCCC2 data set (N = 15,633, M = 360 K) and phenotypes were simulated with M_causal_ = 5000 SNPs explaining *h*^2^_causal_ = 0.2 of phenotypic variance. Reported run times are medians of five identical runs using one core of a 2.27 GHz Intel Xeon L5640 processor. We caution that running time comparisons may vary by a small constant factor as a function of computing environment. FaST-LMM-Select (resp. GCTA-LOCO, EMMAX) memory usage exceeded the 96GB available at N = 15 K (resp. 30 K, 60 K). GEMMA encountered a runtime error (segmentation fault) at N = 30 K. Software versions: FaST-LMM-Select, v2.07; GCTA-LOCO, v1.24; EMMAX, v20120210; GEMMA, v0.94. Numerical data are provided in Supplementary Table 1.

The running time of BOLT-LMM depends not only on the cost of matrix arithmetic, which scales linearly with M and N, but also the number of O(MN)-time iterations required for convergence, which is roughly O(N^0.5^) and also varies with heritability, relatedness, and population structure (see Supplementary Note, Supplementary Fig. 1 and Supplementary Fig. 2). These observations apply both to the full Gaussian mixture modeling performed by BOLT-LMM and to the subset of the computation (steps 1a and 1b) needed to compute BOLT-LMM-inf infinitesimal mixed model association statistics, which in our benchmarks required about 40% of the full BOLT-LMM run time (Fig. 1a and Supplementary Fig. 1a). Our results show that even on very large data sets, BOLT-LMM is efficient enough to enable mixed model analysis using a Gaussian mixture prior, which we recommend because of its potential to increase power.

### Simulations: BOLT-LMM increases power while controlling false positives

To assess the power of BOLT-LMM to detect associated loci, we performed additional simulations using real genotypes from the WTCCC2 data set, which is an ancestry-stratified sample containing both Northern and Southern European samples. We simulated phenotypes with 1250– 10,000 causal SNPs [13,14] explaining 50% of phenotypic variance and an additional 60 candidate causal SNPs explaining 2% of the variance. We further introduced environmental differences in ancestry by including a component of phenotype aligned with the top principal component that explained an additional 1% of the variance. (We note that principal component analysis is not part of BOLT-LMM; our recommendation, consistent with the recommendation of ref. [12], is that it is not necessary to perform PCA when running mixed model association methods.) We chose causal SNPs randomly from the first halves of chromosomes, leaving the second halves of chromosomes to contain only non-causal SNPs (Online Methods).

We computed *χ*^2^ association statistics using linear regression with 10 principal components (PCA) [29], GCTA-LOCO [12], BOLT-LMM-inf, and BOLT-LMM. We were unable to run FaSTLMM-Select [15] because of its memory requirements (Fig. 1). For each method, we computed means of its *χ*^2^ statistics over candidate causal SNPs and compared these means across simulation setups involving different numbers of causal SNPs (Fig. 2a). We observed that BOLT-LMM achieved power gains by modeling non-infinitesimal architectures. For the sparsest genetic architecture (1250 causal SNPs plus 60 causal candidate SNPs), we observed a 25% increase in mean BOLT-LMM *χ*^2^ statistics at candidate SNPs compared to GCTA-LOCO and BOLT-LMM-inf infinitesimal mixed model *χ*^2^ statistics. This metric is readily interpretable as corresponding to a 25% increase in effective sample size; for completeness, we also computed traditional power curves at two significance thresholds (Supplementary Fig. 4). The power gain of the Gaussian mixture model decreased with increasing numbers of causal SNPs (Fig. 2a). This behavior is expected because the advantage of the Gaussian mixture lies in its ability to more accurately model a small fraction of SNPs with larger effects amid a majority of SNPs with near-zero effects. Larger numbers of causal SNPs explaining a fixed proportion of variance result in smaller effect sizes per causal SNP, giving BOLT-LMM less opportunity for power gain. In contrast, all methods other than BOLT-LMM had performance independent of the number of causal SNPs, consistent with the fact that none of these methods model non-infinitesimal genetic architectures. GCTA-LOCO and BOLT-LMM-inf mean *χ*^2^ statistics at candidate causal SNPs were essentially identical and slightly exceeded PCA, consistent with theory [12]. We also tested EMMAX [3] and GEMMA [6], which are vulnerable to proximal contamination [9, 12]; these methods suffered loss of power relative to PCA (Supplementary Fig. 3a), consistent with theory [12].

**Figure 2.**
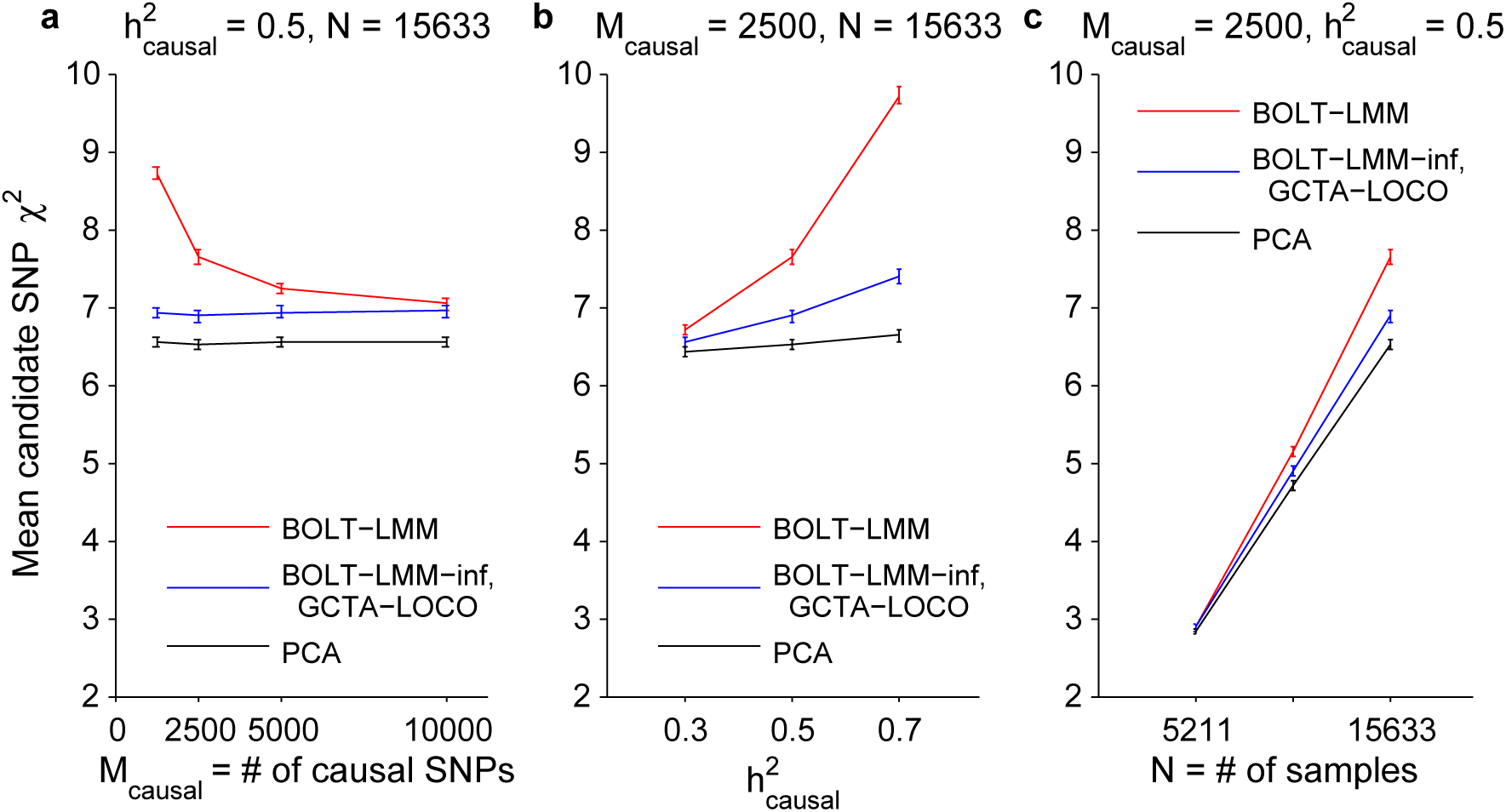
BOLT-LMM increases power to detect associations in simulations. Mean *χ*^2^ at causal candidate SNPs as a function of (**a**) number of causal SNPs, (**b**) proportion of variance explained by causal SNPs, (**c**) number of samples. Simulations used real genotypes from the WTCCC2 data set (N = 15,633, M = 360 K) and simulated phenotypes with the specified number of causal SNPs explaining the specified proportion of phenotypic variance and 60 more candidate SNPs explaining an additional 2% of the variance. Error bars, s.e.m., 100 simulations. We verified on the first 5 simulations that the BOLT-LMM-inf and GCTA-LOCO statistics are nearly identical (Supplementary Table 6). Numerical data are provided in Supplementary Table 2.

To further explore the relationship between the magnitude of Gaussian mixture model power gain and other parameters of the data set, we also varied the proportion of variance explained by causal SNPs (Fig. 2b), and the number of samples in our simulations (Fig. 2c). We observed that the boost in power of BOLT-LMM over infinitesimal mixed model analysis (GCTA-LOCO, BOLT-LMM-inf) increased with each of these parameters. In further simulations using data sets of size N = 30,000 and N = 60,000 (Online Methods) and simulated phenotypes with M_causal_ = 250–15,000 causal SNPs explaining 15–35% of the variance, we observed that the effectiveness of the Gaussian mixture model is closely tied to 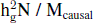 (where 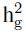 is heritability explained by genotyped SNPs); intuitively, this quantity measures the effective number of samples per causal SNP (Supplementary Fig. 5). These results are consistent with theory, which explains that in the absence of confounding, both infinitesimal and Gaussian mixture model analysis provide a power gain over marginal regression by conditioning on the estimated effects of other SNPs when testing a candidate SNP [9, 12]. As sample size increases, the power gain of both methods approaches an asymptote corresponding to an increase in effective sample size of 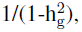 but for sparse genetic architectures, the Gaussian mixture model can approach this asymptote much faster.

To verify that BOLT-LMM is correctly calibrated and robust to confounding, we also computed mean *χ*^2^ statistics across SNPs on the second halves of chromosomes, simulated to all have zero effect (“null SNPs”). Because our simulated phenotypes included an ancestry effect, linear regression without correcting for population stratification suffered 35% inflation. In contrast, the BOLT-LMM and BOLT-LMM-inf statistics were both well-calibrated (Supplementary Fig. 3b, Supplementary Table 3, and Supplementary Table 4). We further verified that Type I error was properly controlled (Online Methods and Supplementary Table 5) and that the distribution of statistics at null SNPs did not deviate noticeably from a chi-square with 1 degree of freedom (Supplementary Fig. 6a,b). Genomic inflation factors [30]) for BOLT-LMM and BOLT-LMM-inf exceeded 1 in these simulations (Supplementary Fig. 6c,d), consistent with polygenicity of the simulated phenotype and use of a mixed model statistic that successfully avoids proximal contamination [12, 13]. In contrast, EMMAX and GEMMA had deflated test statistics (Supplementary Fig. 3b).

Finally,we investigated the similarity between the BOLT-LMM-inf infinitesimal mixed model statistic versus existing methods at the level of individual SNPs. Despite its use of an infinitesimal model, the BOLT-LMM-inf statistic is not identical to any existing mixed model statistic because it is an approximate retrospective test statistic that avoids proximal contamination (Online Methods and Table 1). Nonetheless, we observed that BOLT-LMM-inf statistics very nearly match GCTA-LOCO statistics (which use the standard prospective model), with *R*^2^ > 0.999 (Supplementary Table 6 and Supplementary Fig. 7).

### BOLT-LMM increases power to detect associations for WGHS phenotypes

To assess the efficacy of Gaussian mixture model analysis for increasing power on real phenotypes, we analyzed nine phenotypes in the Women’s Genome Health Study (N = 23,294 samples, M = 324,488 SNPs after QC) (Online Methods). These phenotypes consisted of five lipid phenotypes, height, body mass index, and two blood pressure phenotypes; we chose to analyze these phenotypes because of the availability of published lists of associations from large-scale GWAS of these traits.

We compared the power of three association tests: linear regression with 10 principal components (PCA) [29], infinitesimal mixed model analysis with BOLT-LMM-inf, and Gaussian mixture modeling with BOLT-LMM. Because of memory constraints (Fig. 1), we were unable to run either GCTA-LOCO [12], FaST-LMM [5], or FaST-LMM-Select [15], which are the only previous methods that avoid proximal contamination (Table 1); however, GCTA-LOCO and BOLT-LMM-inf statistics are essentially identical (Supplementary Table 6 and Supplementary Fig. 7). To compare power among these methods, we computed two roughly equivalent metrics: mean *χ*^2^ statistics at known associated loci, a direct but somewhat noisy approach due to having only 19–180 loci for each trait (Supplementary Table 7), and out-of-sample prediction *R*^2^ (measured in cross-validation) using all SNPs for the mixed model methods and using only PCs for linear regression. For mixed model analysis, the latter approach estimates the ability of the mixed model to condition on effects of other SNPs when testing a candidate SNP, which drives its power (Online Methods) [12, 31].

BOLT-LMM achieved higher power than PCA for all traits studied (Fig. 3 and Supplementary Table 8). Most of the increase was due to gains over infinitesimal mixed model analysis, with the magnitude of this power gain increasing with inferred concentration of genetic effects at few loci (Supplementary Table 9). The standard errors of the direct method of assessing improvement (mean *χ*^2^ at known loci) were somewhat high (0.6–2.2%; Fig. 3a and Supplementary Table 8), so the improvement was statistically significant (p < 0.05) for only 6 of the 9 traits. On the other hand, all of the improvements were statistically significant (p < 0.0002) according to the prediction *R*^2^ metric (Fig. 3b and Supplementary Table 8). The largest gains were achieved for lipid traits; for ApoB, a lipoprotein closely related to LDL cholesterol, BOLT-LMM analysis achieved a 10% increase in mean *χ*^2^ statistics versus PCA and a 9% increase versus infinitesimal mixed model analysis at known loci. To verify that these increases were not merely driven by a few loci with the largest effects, we also computed flat averages across loci of improvements in *χ*^2^ statistics (restricting to loci replicating in WGHS with at least nominal *p* < 0.05 significance to reduce statistical noise), obtaining consistent results (Supplementary Table 7). As noted above, simulations show that these improvements will increase with sample size (Fig. 2c and Supplementary Fig. 5).

**Figure 3.**
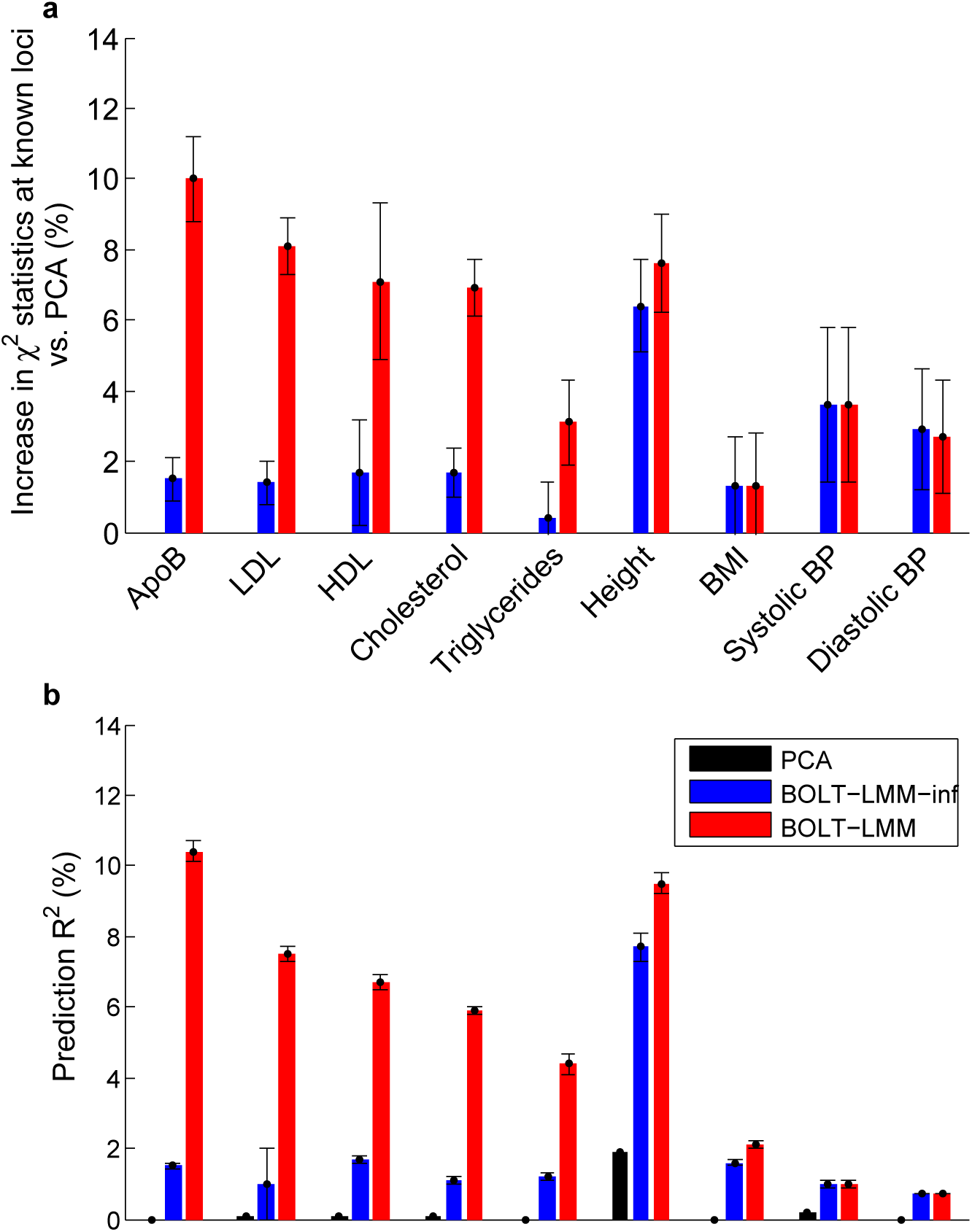
BOLT-LMM increases power to detect associations for WGHS phenotypes. We compare power (measured using two roughly equivalent metrics) of linear regression using 10 principal components, standard (infinitesimal) mixed model analysis, and BOLT-LMM Gaussian mixture model analysis. **(a)** Percent increases in *χ*^2^ statistics across known loci using mixed model methods vs. PCA: ratios of sums of *χ*^2^ statistics over typed SNPs in highest LD with published associated SNPs. **(b)** Prediction *R*^2^ values from 5-fold cross-validation: each fold was left out in turn and predictions were computed by fitting all SNP effects simultaneously (for mixed model methods) or estimating covariate effects (for PCA) using the training folds. The correspondence between association power and prediction accuracy is such that the red bars in **(a)** roughly correspond to differences between red and black bars in **(b)**, and analogously for blue bars (Online Methods). Error bars, jackknife s.e. Numerical data are provided in Supplementary Table 8

We also observed that infinitesimal mixed model analysis achieved statistically significant power gains over PCA, with the magnitude of the power gains increasing with the heritability parameter estimated by BOLT-LMM (Fig. 3 and Supplementary Table 8), which we refer to as pseudo-heritability 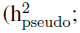 see Online Methods), following ref. [3]. For height 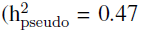 in WGHS), the moderately large sample size of WGHS (N = 23,294) was enough to obtain a 6% increase in BOLT-LMM-inf *χ*^2^ statistics versus PCA, consistent with theory [12, 31]. Once again, larger sample sizes will enable further gains [12, 31].

To verify that BOLT-LMM successfully corrected for confounding from population structure in WGHS, we computed mean *χ*^2^ statistics across all typed SNPs and genomic inflation factors for the three methods compared above as well as uncorrected marginal linear regression. We observed that PCA, BOLT-LMM-inf, and BOLT-LMM statistics were consistently calibrated, while uncorrected linear regression statistics were inflated, especially for height (Supplementary Table 10). We further verified that genetic variation at the lactase gene had a false-positive genome-wide significant association with height using uncorrected marginal regression [32], which disappeared when using PCA, BOLT-LMM-inf, and BOLT-LMM (Supplementary Table 11).

## Discussion

We have described a new algorithm for fast Bayesian mixed model association, BOLT-LMM, and demonstrated that it has time complexity *≈*O(MN^1.5^) and requires only *≈*MN/4 bytes of memory, resulting in orders-of-magnitude improvements in computational efficiency over existing methods for large data sets. We have further shown in simulations and analyses of WGHS phenotypes that the Gaussian mixture modeling capability of BOLT-LMM enables increased association power over standard mixed model analysis while controlling false positives. Among WGHS lipid traits, we observed power increases equivalent to increases in effective sample size of up to 10% over PCA and 9% over standard mixed model analysis.

BOLT-LMM is an advance for two main reasons. First, as sample sizes continue to increase, mixed model analysis is simultaneously becoming more important—in order to correct for population structure and cryptic relatedness in very large data sets—but lees practical with existing methods, all of which have ≥ O(MN^2^) time complexity and high memory requirements. The algorithmic innovations of BOLT-LMM overcome this computational barrier. Second, the ability of BOLT-LMM to better model non-infinitesimal genetic architectures enables a power gain relative to standard mixed model analysis. Recent methodological progress in this direction includes the multi-locus mixed model (MLMM) [7], which identifies and conditions out large-effect loci as fixed effects, and FaST-LMM-Select and related methods [9, 11, 15, 16, 33], which adopt a sparse regression framework that restricts the mixed model to a subset of markers. However, these methods all face the same O(MN^2^) computational hurdle as standard mixed model analysis.

Bayesian methods have previously been developed that apply non-infinitesimal models to improve the accuracy of genetic risk prediction, but translating Bayes factors and posterior inclusion probabilities into the frequentist hypothesis testing framework favored by the GWAS community is a challenge [34]. The variational Bayes spike regression (vBsr) method [35] is a recent step toward addressing this issue, proposing a z-statistic heuristically calibrated by assuming that the vast majority of variants are unassociated (as in genomic control [30]), but such a technique is prone to deflation when large sample sizes cause inflation due to polygenicity [13, 24]. BOLT-LMM sidesteps this difficulty via its hybrid approach of leaving each chromosome out in turn, fitting a Bayesian model on the remaining SNPs, and then applying a retrospective hypothesis test for association of left-out SNPs with the residual phenotype. In contrast to than modeling all SNPs simultaneously and assessing evidence for association using Bayesian posterior inference [34], our approach generalizes existing mixed model methods that are widely used, and we believe its ability to harness the power of Bayesian analysis while still computing frequentist statistics will be useful to GWAS practitioners. Additionally, such a hybrid approach lends itself readily to efficiently testing millions of imputed SNP dosages for association while including only typed SNPs in the mixed model, which we recommend in order to limit computational costs.

While BOLT-LMM improves upon existing mixed model association methods in both speed and power, BOLT-LMM still has limitations. First, the power gain that BOLT-LMM offers over existing methods via its more flexible prior on SNP effect sizes is contingent on the true genetic architecture being sufficiently non-infinitesimal and the sample size being sufficiently large (Supplementary Fig. 5). Second, BOLT-LMM, like existing mixed model methods, is susceptible to loss of power when used to analyze large ascertained case-control data sets in diseases of low prevalence [12]. We recommend BOLT-LMM for randomly ascertained quantitative traits, ascertained case-control studies of diseases with prevalence *≥*5% (e.g., type 2 diabetes, heart disease, common cancers, hypertension, asthma) (see Supplementary Table 12), and studies of rarer diseases in large, non-ascertained population cohorts [36, 37]. For large ascertained case-control studies of rarer diseases, we are developing a method of modeling ascertainment using posterior mean liabilities [38]; applying the techniques of BOLT-LMM to these posterior mean liabilities is an avenue for future research. Third, while mixed model analysis is effective in correcting for many forms of confounding, performing careful data quality control remains critical to avoiding false positives. Fourth, our work does note estimate the heritability explained by genotyped SNPs 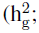 ref. [39])—because 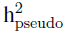 may be different from 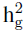 (see Online Methods)—and does not conduct or evaluate genetic prediction in external validation samples from an independent cohort [31]. Fifth, we have not studied the performance of mixed model methods in data sets dominated by family structure [23]; this will be investigated elsewhere. Sixth, while BOLT-LMM extends the infinitesimal model by generalizing the SNP effect prior to a mixture of two Gaussians, other priors are possible and may be more appropriate for some genetic architectures (Table 1 of ref. [19]). Seventh, the running time of BOLT-LMM scales with the number of phenotypes analyzed; for data sets with a very large number of phenotypes (P), the GRAMMAR-Gamma method [10], which has running time O(MN^2^ + MNP) (reviewed in ref. [12]) may be faster. Finally, we have developed fast mixed model analysis for a mixed model with one random genetic effect; extending the algorithm to model multiple variance components [40] is a direction for future work.

**URLs.** BOLT-LMM software, http://www.hsph.harvard.edu/alkes-price/software/.

## Acknowledgments

We are grateful to M. Lipson, S. Simmons, A. Gusev, K. Galinsky, J. Yang, P. Visscher, and Z. Zhu for helpful discussions. This research was funded by NIH grant R01 HG006399 and NIH fellowship F32 HG007805. The WGHS is supported by HL043851 and HL080467 from the National Heart, Lung, and Blood Institute and CA047988 from the National Cancer Institute, the Donald W. Reynolds Foundation and the Fondation Leducq, with collaborative scientific support and funding for genotyping provided by Amgen.

## Online Methods

**Standard mixed model association methods.** Standard methods employ a model

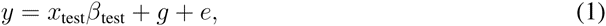

where *y* is the phenotype, *x*_test_ is the candidate SNP being tested, *g* is the genetic effect, and *e* is the environmental effect. Assuming sample size *N*, all of the above are *N*-vectors. We assume for now that all have been mean-centered and there are no covariates; we treat covariates by projecting them out from both genotypes and phenotypes (see Supplementary Note). The genetic and environmental effects are modeled as random effects, while the candidate SNP is modeled as a fixed effect with coefficient *β*_test_, and the goal is to test the null hypothesis *β*_test_ = 0. Under the standard infinitesimal model, the genetic effect is modeled as

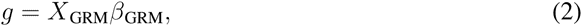

where *X*_GRM_ is an *N × M* _GRM_ matrix, each column of which contains normalized genotypes corresponding to a SNP included in the model, and *β*_GRM_ is an *M* _GRM_-vector of random SNP effect sizes all drawn from the same normal distribution, so that *g* has a multivariate normal distribution with covariance 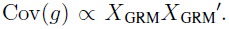 Note that in order to avoid proximal contamination [9, 12], the (*M* _GRM_) SNPs used in *X*_GRM_ should vary depending on which SNP *x*_test_ is being tested: the candidate SNP *x*_test_ (and SNPs in linkage disequilibrium with it) should be excluded from *X*_GRM_ to avoid modeling its effect twice. BOLT-LMM adopts the leave-one-chromosome-out (LOCO) scheme of GCTA-LOCO [12], in which *X*_GRM_ contains all genotyped SNPs except for those on the same chromosome as *x*_test_.

Conventionally, the matrix *X*_GRM_*X*_GRM′_*/M* _GRM_ is referred to as the genetic relationship matrix (GRM) or empirical kinship matrix *K*, and we write

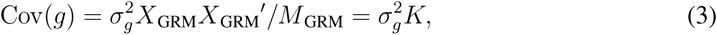

where 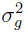 is a variance parameter. Environmental effects are assumed to be i.i.d. normal, so *e* is also multivariate normal with

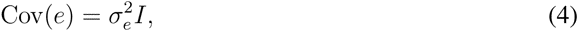

where *I* denotes the *N × N* identity matrix and 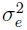 is another variance parameter.

In practice, the variance parameters 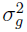 and 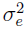 are unknown. Several existing methods [3,10,12] therefore take a two-step approach to computing association statistics: first estimate the variance parameters (with the SNP *x*_test_ removed from the model) using restricted maximum likelihood (REML), and then compute the prospective chi-squared (1 d.o.f.) test statistic (as previously proposed in family-based tests [41])

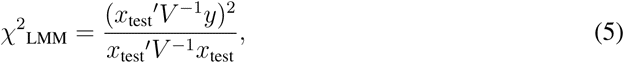

where

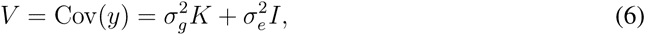

assuming the variance parameters 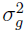 and 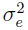 have been fixed to their estimates under the null hypothesis *β*_test_ = 0. Recent computational advances have also enabled computation of exact likelihood ratio test statistics that model the variance parameters while testing the candidate SNP [6,9]. While exact statistics are more accurate in situations with very large-effect SNPs, the approximate methods produce near-identical results in typical human genetics scenarios [3, 10, 12]. In the case of GCTA-LOCO [12], we note that the test statistic becomes

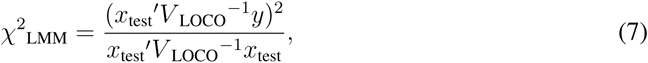

where we have written *V* _LOCO_ for *V* to indicate that the chromosome containing *x*_test_ is left out of the GRM.

**BOLT-LMM-inf mixed model statistic.** The BOLT-LMM-inf infinitesimal mixed model statistic is slightly different:

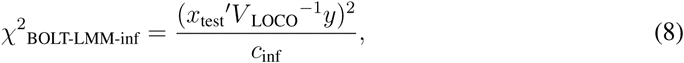

where *c*_inf_ is a constant calibration factor estimated as

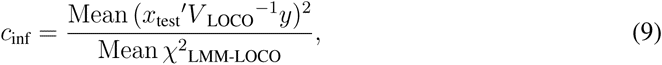

so that

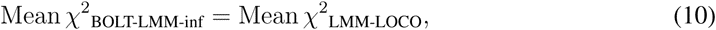

where

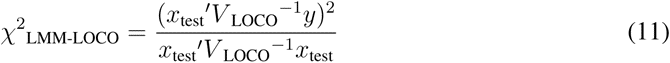

denotes the LOCO version of *χ*^2^_LMM_. In practice, for computational efficiency, we take means over 30 random SNPs not significantly associated with the phenotype (*χ*^2^ < 5 estimated with the GRAMMAR statistic [42]). (For replicability, the BOLT-LMM software uses a pseudo-random number generator to select the “random” SNPs in a deterministic manner so that final results are always exactly the same for different runs.) We have observed empirically in simulations and analyses of real phenotypes that 30 random SNPs are enough to estimate the calibration factor to within 1% (Supplementary Table 13).

We can view the BOLT-LMM-inf statistic either as an approximation of the standard prospective statistic or as a retrospective quasi-likelihood score statistic. The first perspective is motivated by the observation that the denominator of the prospective statistic in equation (5), *x*_test_′*V*^‒1^*x*_test_, is nearly independent of the SNP *x*_test_ being tested [10]. From this perspective, BOLT-LMM-inf is similar to GRAMMAR-Gamma [10], with two key differences: (1) BOLT-LMM-inf is computed via a much faster algorithm (described below) to perform the initial variance components analysis and estimate the calibration constant, and (2) BOLT-LMM-inf avoids proximal contamination via LOCO analysis.

**Retrospective formulation of BOLT-LMM-inf association test.** Alternatively, we can also view BOLT-LMM-inf as a retrospective quasi-likelihood score test similar to *T ^SCORE-R^* [43] and MASTOR [23]. Following ref. [23], this viewpoint models the phenotype *y* as known and the SNP *x*_test_ as random, being drawn from a distribution with covariance Cov(*x*_test_) = Φ and mean

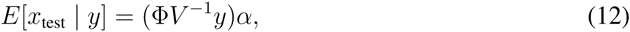

where we wish to test the null hypothesis *α* = 0. Note that *x*_test_ is a discrete random variable, so we cannot formally model it as a multivariate normal and perform likelihood-based analysis, but the retrospective mean model is enough to obtain a quasilikelihood score test statistic that is asymptotically *χ*^2^-distributed:

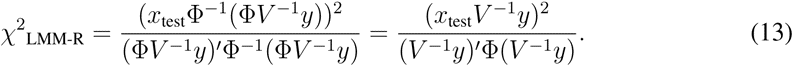

Previous formulations of this statistic [23, 43] have been aimed at family studies in which the pedigree matrix may be used as the covariance matrix Φ. For the general case of individuals with unknown pedigree, a seemingly natural choice is to substitute the GRM, but unfortunately, the GRM does not accurately reflect the covariance matrix of the distribution of unassociated (i.e., null) SNPs. In particular, for large sample sizes, this approach incurs deflation because it incorporates SNPs associated with *y* into the covariance model (for *x*_test_, treated as random), causing overestimation of the variance of *x*_test_*V ^-^*^1^*y*. This phenomenon is not overcome by LOCO analysis because unlike with prospective modeling, the problem is not proximal contamination; rather, it is polygenicity: to obtain unbiased variance estimates based on genotype data, we would need to eliminate associated SNPs, which runs into the same difficulty faced by genomic control [13, 24].

While the denominator of *χ*^2^_LMM-R_ in equation (13), (*V*^‒1^*y*)′Φ(*V*^‒1^*y*), cannot be computed without Φ, it does not involve the candidate SNP *x*_test_ and can therefore be treated as a constant calibration factor, so that in practice, equation (13) is equivalent to the BOLT-LMM-inf formulation in equation (8). (More precisely, within a LOCO scheme, the matrix *V* depends on which chromosome *x*_test_ belongs to, so there should formally be a different denominator for each chromosome, but all are approximately equal.) The retrospective model thus provides an alternative justification for the BOLT-LMM-inf statistic.

**BOLT-LMM Gaussian mixture model association statistic.** Importantly, the retrospective viewpoint also enables a natural generalization of BOLT-LMM-inf to a more powerful association test for non-infinitesimal phenotypes. The basic observation motivating this extension is that in the retrospective mixed model statistic *χ*^2^_LMM-R_, the vector *V ^-^*^1^*y* is a scalar multiple of the residual after best linear unbiased prediction (BLUP)—i.e., the component of the phenotype remaining after conditioning out SNP effects estimated by the mixed model, which drives the power of mixed model association—but this vector may be replaced by any other vector not involving *x*_test_. We therefore define a generalized statistic

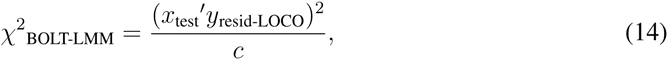

where *y*_resid-LOCO_ denotes the residual phenotype upon conditioning out the effects of SNPs not on the same chromosome as *x*_test_, estimated by fitting a Gaussian mixture extension of the standard LMM, and *c* denotes a constant calibration factor, estimated so that the LD Score regression intercept [24] of *χ*^2^_BOLT-LMM_ matches that of the calibrated *χ*^2^_BOLT-LMM-inf_ statistic. When using the infinitesimal model, *y*_resid-LOCO_ is proportional to *V ^-^*^1^*y*, and *χ*^2^_BOLT-LMM_ reduces to *χ*^2^_BOLT-LMM-inf_.

The Gaussian mixture extension of the standard LMM is defined as follows. First, note that the null model of the standard LMM used by BOLT-LMM-inf admits the Bayesian formulation

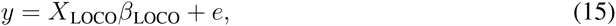

where SNP effects *β_m_* (where *m* indexes SNPs not on the left-out chromosome) are independently drawn from the normal distribution

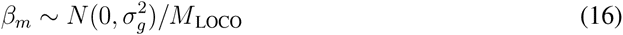

and environmental effects *e_n_* (where *n* indexes samples) are independently drawn from the normal distribution 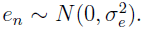 Performing best linear unbiased prediction amounts to computing the posterior mean of the genetic effect *X*_LOCO_*β*_LOCO_.

To generalize this model to non-infinitesimal genetic architectures, we replace the normal prior on SNP effect sizes with a spike-and-slab mixture of normals [19]:

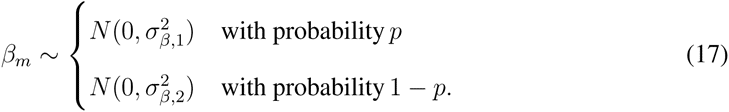

The rationale behind using a mixture of normals is that it more flexibly models the heavier-tailed distributions of genetic effects of typical (non-infinitesimal) phenotypes. Explicitly, if *p* ≪ 1 and 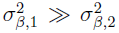 small, the first component of the mixture is a “slab” that models the existence of a small number of relatively large-effect loci, while the second component is a “spike” that models the assumption that most SNPs have near-zero—but not exactly zero—effect on the phenotype. It is important that the second component not be a pure spike with zero variance, because assigning a small amount of variance to the spike component allows the model to capture components of phenotype explained by aggregate genome-wide effects such as ancestry or relatedness. Consequently, when testing SNPs for association with the residual phenotype, these genome-wide effects are appropriately conditioned out, making BOLT-LMM robust to confounding.

BOLT-LMM estimates best-fit mixture parameters to use in the Gaussian mixture model via cross-validation; details are given in the algorithm description below. We can now finally complete the definition of the *χ*^2^_BOLT-LMM_ statistic in equation (14): using the fitted mixture parameters, we fit the Bayesian model (once per left-out chromosome) and obtain residuals

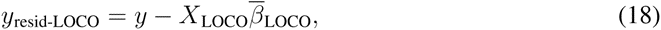

where 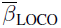 are the estimated posterior mean effect sizes. Plugging these residuals into equation (14) gives the BOLT-LMM Gaussian mixture model association test statistic.

**Fast iterative algorithm.** As described in Results (Overview of Methods), the BOLT-LMM software performs a four-step computation for mixed model association analysis, stopping after the first two steps when specialized to the infinitesimal model. We outline the main algorithms here and provide full details in the Supplementary Note.

**Step 1a: Estimate variance parameters.** A key advance of BOLT-LMM is its ability to perform REML variance components analysis (for the special case of one identity and one non-identity variance component)—i.e., estimate 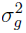 and 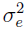 in equations (3) and (4)—using linear-time iterations without building or decomposing any covariance matrices. Our algorithm relies on rewriting or approximating all O(MN^2^)-time matrix computations in terms of solutions to linear systems of equations (i.e., expressions of the form *V ^-^*^1^*x*), which we in turn compute using only O(MN)-time matrix-vector products via conjugate gradient iteration. In particular, computing *V ^-^*^1^*x* using conjugate gradient iteration only requires computing products of *V* with vectors, which we can compute in O(MN)-time without forming *V* by leaving *V* in factored form 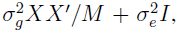 distributing the product, and multiplying from right to left.

To estimate variance parameters using this approach, we apply the observation that the REML first-order conditions on 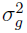 and 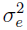 are equivalent to the system of equations

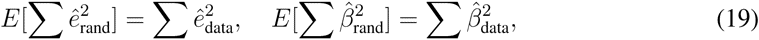

where both the left and right sides of each equation are functions of the assumed variance parameters 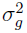 and 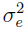 (equation (14.8) of ref. [44]). The right sides involve BLUP predictions on the observed phenotype data *y* (performing BLUP using the assumed 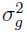 and 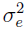 which can be done using conjugate gradient iteration), while the left sides are expectations of the same quantities *over random y generated according to the model y* = *g* + *e with variance parameters set to the assumed* 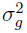 *and* 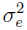 We can therefore estimate the left sides (for a particular choice of 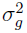 and 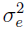) by Monte Carlo sampling, i.e., by generating random *y*_rand_ = *Xβ*_rand_ + *e*_rand_ from 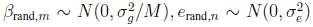 and then performing BLUP on *y*_rand_ using the assumed 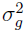 and 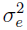 Only a few Monte Carlo trials are required; BOLT-LMM computes 3–15 trials, with the number of trials decreasing as *N* increases, reflecting the way in which the standard error depends on sample size and the number of trials. Explicitly, BOLT-LMM by default uses 4 *×* 10^9^*/N* ^2^ trials, with a minimum of 3 and a maximum of 15. The number of trials can also be varied as a user-specified parameter.

While the above procedure as stated requires a 2-parameter search over 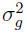 and 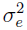 we can eliminate 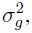 leaving a 1-parameter zero-finding problem over the parameter 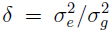 [45]. BOLT-LMM performs the zero-finding numerically using secant iteration, a finite difference approximation of Newton’s method. Finally, we note that satisfying the first-order conditions is not in general a sufficient condition to have found the REML global optimum, but for the case of only one non-identity variance component, the likelihood surface appears to be well-behaved, and we have never observed an instance in which this estimation procedure failed.

**Step 1b: Compute and calibrate BOLT-LMM-inf statistics.** Using the variance parameters estimated in step 1a, it is straightforward to compute (for each LOCO rep) the quantity *V* _LOCO_*^-^*^1^*y* in the numerator of the BOLT-LMM-inf statistic, equation (8), by using conjugate gradient iteration as described above. Completing the computation of the numerator for each SNP *x*_test_ then just amounts to a calculating a dot product, which requires only O(MN) additional cost across all SNPs. Similarly, to compute the calibration constant *c*_inf_ in equation (9), BOLT-LMM rapidly computes the prospective statistic *χ*^2^_LMM-LOCO_ from equation (11) at 30 random SNPs by applying conjugate gradient iteration to compute *V* _LOCO_*^-^*^1^*x*_test_ for each of the 30 selected SNPs *x*_test_.

Note that there is a slight mismatch between the variance parameters computed in step 1a, which BOLT-LMM computes once using all SNPs—not leaving any chromosomes out—and the theoretically optimal variance parameters for the infinitesimal mixed model fit in each LOCO rep, which vary slightly depending on which chromosome is left out. However, we have observed in simulations that slight mis-specification of the variance parameters has a negligible impact (*<*0.5%) on the calibration of the BOLT-LMM-inf and BOLT-LMM statistics (Supplementary Table 4). Because very slight miscalibration is not a concern for confounding from population stratification at highly differentiated markers (Supplementary Table 11) and has little impact on Type I error (Supplementary Table 5), the BOLT-LMM software does not by default refit variance parameters for each LOCO rep. If extremely precise calibration is desired, we provide runtime options to refit variance parameters for each LOCO rep or to partition the genome more finely (e.g., into 100 segments rather than 22), each at the cost of a factor of 2–3 in running time.

**Step 2a: Estimate Gaussian mixture prior parameters.** The first step of BOLT-LMM Gaussian mixture model association analyis is to estimate parameters for the mixture of two Gaussians used as the Bayesian prior on SNP effect sizes. As written in equation (17), this mixture has three parameters: 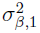 and 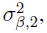 the variances of the two Gaussians, and *p*, the probability of drawing from the first Gaussian. To reduce the complexity of parameter estimation, we constrain the total variance of the mixture to equal the variance 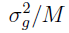 estimated for SNP effects by the infinitesimal model in step 1a:

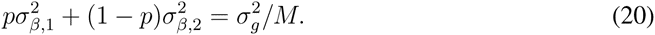

We reparameterize the remaining two degrees of freedom using the parameters *p* and *f*_2_, where *f*_2_ denotes the proportion of the total mixture variance within the second Gaussian (the “spike” component that models small genome-wide effects):

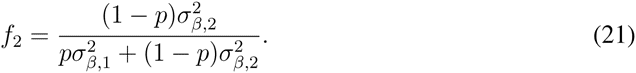

Because the model fit is not sensitive to the precise values of the mixture parameters, we test a discrete set of model parameter combinations:

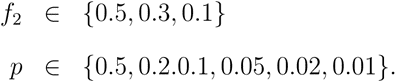

Note that *f*_2_ = 0.5*, p* = 0.5 corresponds to the infinitesimal model: when *f*_2_ = 1 *-p*, the two Gaussians are identical and the mixture is degenerate. We bounded *f*_2_ from below to ensure that at least a small amount (10%) of the mixture variance is assigned to the spike component, protecting against confounding from genome-wide effects. We bounded *p* from below to prevent the model from trying to fit too strongly to a few SNPs, which makes Bayesian analysis computationally difficult and also increases susceptibility to confounding.

BOLT-LMM performs model selection among the 18 possible parameter pairs (*f*_2_*, p*) by performing cross-validation to optimize mean-squared prediction *R*^2^. More precisely, for each parameter pair, for each cross-validation fold, BOLT-LMM estimates posterior mean SNP effect sizes *β_m_* on the training data (using a variational Bayes procedure, described in the next paragraph, to fit the model) and uses these effect size estimates to make predictions on the left-out fold, which are then compared to the actual left-out phenotype values. The parameter pair (*f*_2_*, p*) producing the highest prediction *R*^2^ (mean across folds) is then selected. However, if the best model only reduces prediction error by less than 1% (in absolute prediction *R*^2^) compared to the infinitesimal model, BOLT-LMM terminates the Gaussin mixture analysis and does not proceed to step 2b to compute association statistics. We implement this feature as the default option (which may be overriden) to save unnecessary computation in situations where the Gaussian mixture model provides little or no improvement in association power.

BOLT-LMM uses a variational approximation algorithm to fit Bayesian linear regressions with Gaussian mixture priors. Approximation methods are necessary for Bayesian inference in this setting because analytic formulas for exact posterior means involve intractable integrals. We apply a fully factored variational approximation [21, 22, 35] that repeatedly loops through the SNPs, updating the estimated effect size of each SNP with its posterior mean conditional on current estimates of all other SNP effects. Explicitly, we begin by initializing each SNP effect *β_m_* to 0 and initializing the residual phenotype *y*_resid_ = *y − Xβ* to *y*. Then, each iteration performs the following loop over SNPs *m* = 1*,…, M*:

1. Remove effect of SNP *m* from residual. *y*_resid_*_,−m_ ← y*_resid_ + *x_m_β_m_*
2. Re-estimate effect of SNP *m*. *β_m_←* posterior mean given *y*_resid_*_,−m_*
3. Replace effect of SNP *m* in residual. *y*_resid_ *← y*_resid_*_,−m_ − x_m_β_m_*

This iteration has also previously been termed “iterative conditional expectation (ICE)” [20]. The variational Bayes framework puts this iteration on a sound theoretical footing as an optimization of an approximate log likelihood function; the iteration monotonically increases this function and is guaranteed to converge [46]. In fact, we show that the optimization can be reformulated as cyclic coordinate descent applied to a penalized regression problem arising from Bayesian linear regression using a transformed prior (Supplementary Note). The approximate log likelihood also serves as a convenient convergence criterion: BOLT-LMM stops the iteration when the absolute improvement in approximate log likelihood from one update cycle through all the SNPs drops below 0.01.

One difference between the algorithm BOLT-LMM applies and previous variational approaches to genetic modeling is that we do not estimate model hyperparameters within the variational iteration [22, 35] or based on variational approximate log likelihoods [21]. Instead, BOLT-LMM uses cross-validation [15], which we found to be more robust to decreases in the quality of the variational approximation caused by linkage disequilibrium.

**Step 2b: Compute and calibrate BOLT-LMM Gaussian mixture model statistics.** After inferring parameters of the mixture prior in step 2a, BOLT-LMM uses the same variational Bayes algorithm to estimate posterior mean residuals *y*_resid-LOCO_ (independently for each left-out chromosome). The numerators of the BOLT-LMM Gaussian mixture model statistic given in equation (14) are then easily obtained as dot products with test SNPs, leaving only the constant calibration factor *c* in the denominator to be calculated. Unlike in the case of the infinitesimal model, here we do not have a prospective statistic to calibrate against, so we instead apply LD Score regression [24]. LD Score regression observes that for complex traits, properly calibrated *χ*^2^ association statistics approximately obey the following linear trend (on average across the genome):

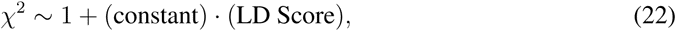

where loosely speaking, the LD Score of the test SNP is its effective number of LD partners. Because the intercept of this regression relation is 1, it follows that if we have computed a set of *χ*^2^ statistics up to an unknown constant factor *c*, then one way to estimate *c* is to compute the intercept of the regression of *χ*^2^ statistics on LD Scores.

To achieve very precise calibration, BOLT-LMM adopts a slightly more complex procedure that matches the intercept of LD Score regression on *χ*^2^_BOLT-LMM_ (after calibration) to that of the properly-calibrated infinitesimal statistic *χ*^2^_BOLT-LMM-inf_. We do so to increase robustness of the calibration to violations of the modeling assumptions underlying LD Score regression, which may result in attenuation bias (i.e., inflation of the intercept toward the mean *χ*^2^) that causes the straightforward LD Score correction to association statistics to be conservative, though still much less conservative than genomic control. Because LD Score regression on either *χ*^2^_BOLT-LMM_ or *χ*^2^_BOLT-LMM-inf_ should be affected roughly equally by attenuation bias, matching LD intercepts guards against this potential problem.

**Simulations.** We simulated data sets from the WTCCC2 data with the goal of creating realistic test scenarios including linkage disequilibrium and possible confounding from population stratification or relatedness. The WTCCC2 data set contains 15,633 samples (10,204 cases and 5,429 controls) genotyped at 360,557 SNPs after QC; full details are given in ref. [12]. We simulated genotype data with N = 3,750 to 480,000 individuals and M = 150,000 or 300,000 SNPs by subsampling the desired number of SNPs from the WTCCC2 data set and then independently building N individuals as mosaics of individuals from the original data set using the following procedure. First, specify a number of “ancestors” *A* for each individual to have. Then, for each simulated individual, select *A* ancestors at random from among the original individuals and create the simulated individual’s genotype data by chopping the genome into segments of 1000 SNPs and copying each segment from a randomly sampled (diploid) ancestor.

Note that this approach retains realistic LD among SNPs, and for small values of *A*, it also retains population structure and introduces relatedness among individuals that share ancestors. In our simulations, we used *A* = 2 (substantial structure and relatedness) or *A* = 10 (low levels of structure and relatedness).

We simulated phenotypes as sums of up to four components: genetic effects from a specified number of causal SNPs, genetic effects from a specified number of candidate causal SNPs, population stratification, and environmental effects. We simulated genetic effects by selecting SNPs at random from the first half of the list of typed SNPs in each chromosome and at least 2Mb and 2cM before the middle SNP. (Likewise, when assessing calibration at null SNPs, we only tested SNPs in the second halves of chromosomes and at least 2Mb and 2cM beyond the middle SNP, to avoid contamination of null SNPs from LD with causal SNPs.) We sampled effect sizes for the selected normalized SNPs from a standard normal distribution. We simulated population stratification by taking the first principal component of the normalized genotype matrix. We sampled environmental effects for all individuals from a standard normal. To create phenotypes with specified proportions of variance explained by each of these effects, we normalized each component (an *N*-vector across samples) and formed weighted sums using the desired weights.

Because causal SNPs in our simulated phenotypes are selected only from among genotyped SNPs, we computed reference LD Scores (used to calibrate the BOLT-LMM statistic in our simulations) by summing LD only to typed SNPs—as opposed to all SNPs, which is appropriate for real phenotypes [24]—in 379 European-ancestry samples from the 1000 Genomes Project [47].

**WGHS data set.** The Women’s Genome Health Study (WGHS) is a prospective cohort of initially healthy, female North American health care professionals at least 45 years old at baseline representing participants in the Women’s Health Study (WHS) who provided a blood sample at baseline and consent for blood-based analyses. The WHS was a 2 × 2 trial beginning in 1992-1994 of vitamin E and low dose aspirin in prevention of cancer and cardiovascular disease with about 10 years of follow-up. Since the end of the trial, follow-up has continued in observational mode. Genotyping in the WGHS sample was performed using the HumanHap300 Duo “+” chips or the combination of the HumanHap300 Duo and iSelect chips (Illumina, San Diego, CA) with the Infinium II protocol. For quality control, all samples were required to have successful genotyping using the BeadStudio v. 3.3 software (Illumina, San Diego, CA) for at least 98% of the SNPs. A subset of 23,294 individuals were identified with self-reported European ancestry that could be verified on the basis of multidimensional scaling analysis of identity by state using 1443 ancestry informative markers in PLINK v. 1.06. In the final data set of these individuals, a total of 339,596 SNPs were retained with MAF *>*1%, successful genotyping in 90% of the subjects, and deviations from Hardy-Weinberg equilibrium not exceeding *p* = 10*^-^*^6^ in significance. In our analyses, we further eliminated non-autosomal SNPs, duplicate SNPs, and custom SNPs, leaving 324,488 SNPs.

**Correspondence between association power and prediction accuracy.** Intuitively, mixed model analysis to test a candidate marker for association gains power over linear regression (assuming no confounding) by conditioning on other markers modeled as random effects. This intuition is especially clear in the BOLT-LMM statistical formulation, where we retrospectively test candidate markers for association with residual phenotypes. Residualizing eliminates the component of phenotype that was successfully predicted by other markers, so that the subsequent association test is trying to detect candidate marker association signal amid phenotypic variance (“noise”) that has been decreased by a factor of 1-*R*^2^. Note that residualizing leaves the amount of candidate marker signal unchanged, assuming no proximal contamination and assuming a randomly ascertained phenotype; violation of these assumptions results in loss of power [12].

Quantitatively, a reduction of association test noise by a factor of 1-*R*^2^ results in an increase in *χ*^2^ statistics by a factor of 1/(1-*R*^2^) at truly associated loci. (More precisely, the quantity *χ*^2^-1 increases by 1/(1-*R*^2^) on average, but for *χ*^2^ ≫ 1 at known loci, this distinction is minor.) This relationship can be seen explicitly in comparing equation (1) of ref. [31] for prediction *R*^2^ using BLUP,

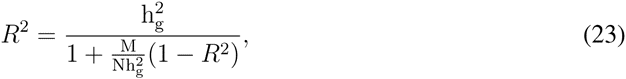

with Supplementary Table 2 of ref. [12], which gives the following formulas for mean *χ*^2^ statistics using linear regression vs. mixed model association:

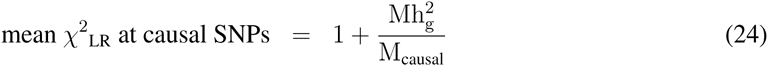

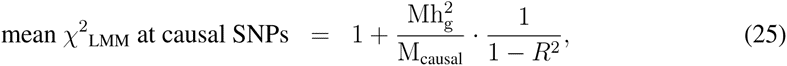

where *R*^2^ in equation (25) (denoted 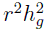 in ref. [12]) satisfies the same quadratic equation (23).

In Fig. 3a, we plot increases in mean *χ*^2^ statistics at known loci for mixed model methods vs. PCA, and in Fig. 3b, we plot absolute prediction *R*^2^. As explained above, the increase in mean *χ*^2^ statistics at known loci for mixed model methods vs. linear regression—assuming no confounding—is approximately 1*/*(1 *-R*^2^) *-* 1 *≈ R*^2^. When including principal components in linear regression to avoid confounding, we get an increase in mean *χ*^2^ statistics at known loci for mixed model methods vs. PCA of approximately *R*^2^_LMM_ *-R*^2^_PCA_, where the latter is the prediction *R*^2^ obtained by linear prediction using only the principal components. Thus, the red bars in Fig. 3a roughly correspond to differences between red and black bars in Fig. 3b, and analogously for blue bars. The correspondence is approximate because of the approximations mentioned above and two additional subtleties: (1) the LOCO scheme that BOLT-LMM uses to avoid proximal contamination renders the mixed model unable to condition out effects of markers on the same chromosome as the candidate marker; (2) our measurement of prediction *R*^2^ uses 5-fold cross-validation, reducing the training sample size by 20%. However, these two effects have small magnitudes and act in opposite directions, so that ultimately the correspondence is very tight, as is visually apparent in Fig. 3.

**Pseudo-heritability.** Following ref. [3], we refer to the heritability parameter inferred from variance components analysis as “pseudo-heritability,” denoted 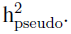 Note that 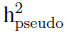 can be larger than 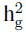 (the heritability explained by genotyped SNPs [39]) due to cryptic relatedness or population structure [48], but for samples such as WGHS without substantial relatedness or population substructure, 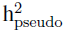 is close to 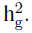

## Supplementary Note

### 1 BOLT-LMM algorithm details

The BOLT-LMM algorithm is overviewed in Online Methods. Here, we provide additional details for components of the computation not fully described earlier.

#### 1.1 Model setup

##### 1.1.1 Initialization: Missing data, normalization, and covariates

Given a data set of genotypes and phenotypes, we apply the following procedure to create a normalized genotype matrix *X*. We begin by dealing with missing data as follows. First, we perform QC by filtering out SNPs and individuals with missing rates exceeding thresholds (default 10% for each). Second, we filter out individuals with missing phenotypes. Third, if the analysis includes covariates, we filter out individuals with any missing covariates. (As an alternative to this approach, known as “complete case analysis,” we also implement the option to use the “missing indicator method,” which adds missing status indicator variables as additional covariates.) Finally, we replace missing genotypes in the remaining data with per-SNP averages. We denote by *N* and *M* the numbers of samples and SNPs remaining post-QC.

Next, we mean-center each SNP and normalize the SNPs to have equal sample variance. We also mean-center the phenotypes. We model covariates by projecting them out from both genotypes and phenotypes, which is equivalent to including them as fixed effects. Explicitly, we compute a basis spanning the covariate vectors and subtract out the components of the SNP and phenotype vectors along the basis vectors. This procedure is mathematically equivalent to multiplying by an orthogonal projection matrix **P**_fixed_ that projects vectors to an (*N − C*)-dimensional subspace 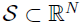 orthogonal to the covariates, where *C* is the number of independent covariates (including the all-1s vector, which is implicitly included as a covariate upon mean-centering). For comparison, GCTA-LOCO [12] version 1.24 provides two options for treating covariates: (1) projecting out covariates from the phenotype vector only (the default, which is an approximate approach); or (2) fitting covariates as fixed effects along with the candidate SNP in each association test (the `--mlma-no-adj-covar` option, which is equivalent to projecting covariates out from both genotypes and phenotypes). The GCTA-LOCO documentation notes that the latter option significantly reduces the computational efficiency of GCTA-LOCO, but the BOLT-LMM implementation is not subject to this loss of computational efficiency.

We denote by *X* the final *N × M* matrix of genotypes and *y* the final *N*-vector of phenotypes after QC, normalization, and projection. While *y* and all columns of *X* (i.e., SNP vectors) are *N*-dimensional vectors, it is important to keep in mind that they all actually belong to the (*N − C*)-dimensional subspace *S* left after projecting by **P**_fixed_. For example, when estimating variance parameters, accounting for the loss of degrees of freedom distinguishes restricted maximum likelihood (REML) analysis—the preferred approach, which we implement—from maximum likelihood (ML), and when computing *χ*^2^ test statistics, we need to use *N − C* instead of *N* as the sample size.

##### 1.1.2 Models

We therefore use the following general model to estimate hyperparameters (i.e., variance parameters under the infinitesimal model and Gaussian mixture parameters under the non-infinitesimal model):

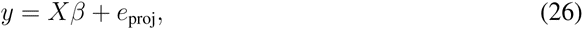

where

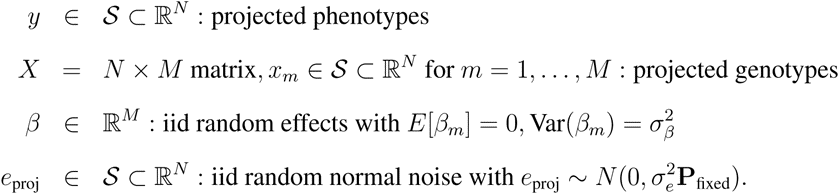

Reiterating the point above, this “mixed” model does not contain any fixed effects; the fixed effects have been projected out, so that the entire model—including the noise term *e*_proj_—lives in the subspace *S*.

**Model for estimating infinitesimal model parameters.** When estimating variance parameters under the infinitesimal model in Step 1a of the BOLT-LMM algorithm, we assume the SNP effect prior is the normal distribution

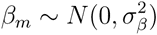

and we estimate 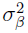 and 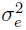 using the stochastic REML approximation algorithm described in Section 1.2. Note that the following notation is often used for the total genetic effect *Xβ* and its covariance:

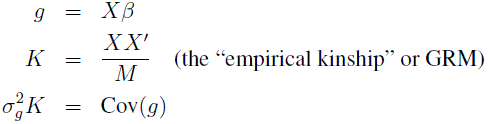

in which case

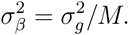

**Model for estimating Gaussian mixture model parameters.** When estimating Gaussian mixture parameters in Step 2a of the BOLT-LMM algorithm (after having obtained estimates of 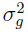 and 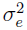 in Step 1a), we generalize the SNP effect prior to the three-parameter mixture of normals

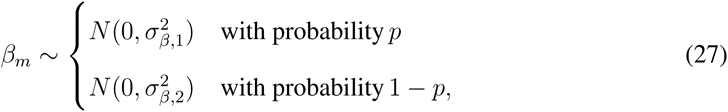

where we require

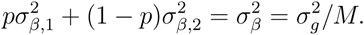

We reparameterize the remaining two degrees of freedom using the parameters *f*_2_ and *p*, where *f*_2_ denotes the proportion of the total mixture variance within the second Gaussian (the “spike” component that models small genome-wide effects):

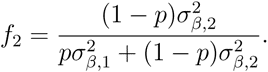

Thus, Step 2a consists of estimating *f*_2_ and *p*.

**Models for computing association statistics.** When performing association tests in Steps 1b and 2b of the BOLT-LMM algorithm, we modify the above models slightly, including the SNP being tested as a fixed effect, and leaving out all SNPs on its chromosome from the random effect:

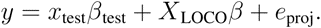

As discussed in Online Methods, we do not by default re-estimate the hyperparameters 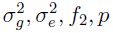 when performing association tests; instead, we simply reuse the estimates obtained above from the (slightly different) models above, which include all SNPs in the random effect (and no fixed effect). However, we also offer the option of re-estimating 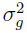 and 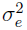 with each chromosome left out in turn.

#### 1.2 Variance component estimation: Stochastic REML approximation

Here we provide details of our stochastic algorithm for estimating REML variance parameters in Step 1a of BOLT-LMM. As described in Online Methods, the crux of the method is to employ the observation that the REML first-order conditions on 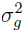 and 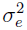 are equivalent to the system of equations

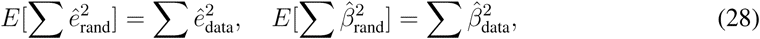

where both the left and right sides of each equation are functions of the assumed variance parameters 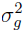 and 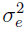 [44, equation (14.8)]. (The summations are over components of each vector, which is a slight abuse of notation for 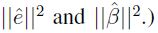 On the right sides, 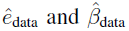 are best linear unbiased predictions (BLUP) on the observed phenotype data assuming the basic linear mixed model *y* = *Xβ* + *e*_proj_, equation (26), with

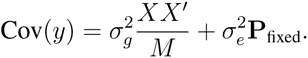

On the left sides, expectations are taken over the same quantities, with the phenotype data replaced by *random y*_rand_ *generated according to the model y* = *Xβ* + *e*_proj_ with variance parameters set to the assumed 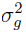 and 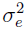.

Explicitly, defining

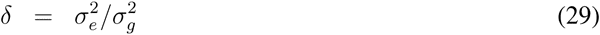

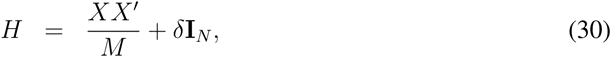

the BLUP estimates 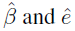 for a phenotype vector *y* are:

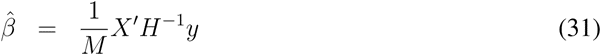

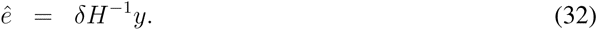

(Note that technically, the scaled covariance matrix is actually 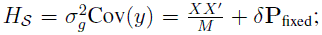 however, because all of our vectors belong to the subspace *S*, using **I***_N_* instead of **P**_fixed_ produces the correct result and has the convenience of making the operator *H* invertible in the full space 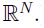)

Equations (29)–(32) show that for a fixed value of *δ*, the BLUP predictions 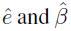 are constant. Thus, the right sides of the REML first order conditions (28), involving BLUP on the observed phenotypes, depend only on the variance ratio *δ* and are independent of the variance scale, which we may parameterize by 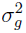 (in which case 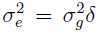 becomes a dependent variable). The left sides scale proportionately with 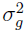 because scaling up the variances scales up randomly generated phenotypes *y*_rand_. Therefore, finding 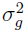 and 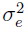 that solve the pair of equations (28) is equivalent to: (1) finding the value of the single parameter *δ* such that

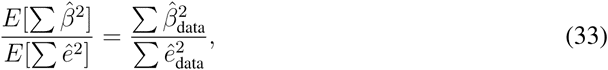

and (2) choosing 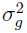 to scale the expectations to match the values observed on the data. (This procedure is analogous to the usual REML trick of optimizing over *δ* and then setting 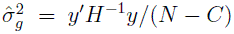 [45].)

We propose an algorithm that rapidly estimates the ratio of expectations in equation (33) and uses this estimate within a one-parameter search over *δ*. Define

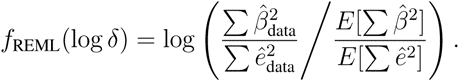

For a fixed value of *δ*, we produce 3–15 (decreasing with *N)* Monte Carlo estimates of the expectations by generating random phenotypes

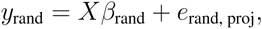

where

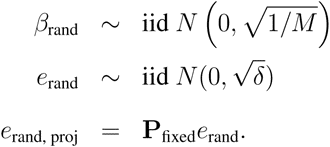

(Note that as the variance scale parameter 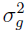 is irrelevant for this part of the computation, we set it to 1 for convenience. Note also that the use of the projection matrix **P**_fixed_ is what makes this procedure compute REML rather than ML estimates.) We then run BLUP (using the chosen value of *δ*) using conjugate gradient iteration [26, 27] to obtain Monte Carlo estimates of 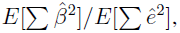 and we likewise run BLUP to compute 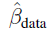 and *ê*_data_, which together give an estimate 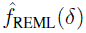 of *f*_REML_(*δ*).

We wish to find *δ* such that *f*_REML_(log *δ*) = 0. We have observed empirically in simulations and real data sets that *f*_REML_(log *δ*) increases monotonically with *δ* except possibly at extremely small or large values of *δ* outside the range of reasonable parameter values (corresponding to, say, 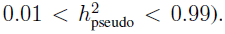 Thus, one approach to finding the unique zero of *f*_REML_ within the reasonable parameter range is binary search, with the caveat that some care is needed because we can only compute noisy Monte Carlo estimates 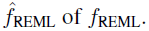

We implement the following more robust approach. Instead of independently re-randomizing phenotypes *y*_rand_ used to compute estimates 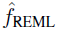 at different values of *δ*, we a generate a single set of random phenotypic component pairs

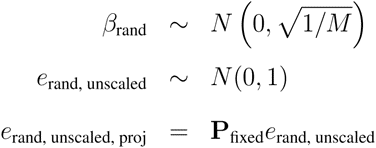

and then use these component pairs to generate random phenotypes

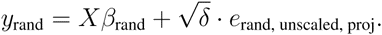

for any given *δ*. Using this approach, the Monte Carlo estimate 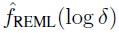 becomes a smooth function of log *δ*, allowing us to use the secant method, a finite difference approximation of Newton’s method, to perform zero-finding on 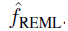

#### 1.3 Gaussian mixture model fitting: Variational Bayes iteration

As outlined in Online Methods, the core component of the BOLT-LMM Gaussian mixture modeling algorithm is a variational iteration [20–22, 35] that computes approximate posterior mean effect sizes *β_m_* for the Gaussian mixture model version of equation (26). We sketch the iteration in slightly more detail here, leaving a full description for Section 2.

The idea of the iteration is to obtain successively better approximations of the SNP effect sizes by cyclically updating each estimated SNP effect with its posterior mean conditional on the current estimates of all other SNP effects. Explicitly, we begin by initializing each SNP effect *β_m_* to 0 and initializing the residual phenotype *y*_resid_ = *y − X β* to *y*. Then, each iteration performs the following loop over SNPs *m* = 1*,…, M*:

Step 2 amounts to computing the posterior mean of *β_m_* given its assumed mixture prior in equation (27), the environmental noise distribution 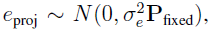 and the residual phenotype *y*_resid_*_,−m_* given current estimates of all other SNP effects. Steps 1 and 3 are a computational technique allowing the residual in Step 2 not to be computed from scratch for each SNP, so that an entire iteration (updating each SNP effect once) takes *O*(*M N)* time.

The basic structure of this iteration is the shared with previous variational methods [20–22,35], with the update Step 2 varying among methods depending on the assumed SNP effect prior. As noted in Online Methods, an additional slight difference is that BOLT-LMM estimates the hyperparameters 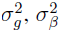 using REML and *f*_2_, *p* using cross-validation [15], whereas previous approaches have performed hyperparameter estimation within the variational iteration [22, 35] or using the variational approximate log likelihood [21]. We used cross-validation instead because we found cross-validation estimates to be more robust to decreases in the quality of the variational approximation caused by linkage disequilibrium.

#### 1.4 Cross-validation for estimating Gaussian mixture parameters

We perform 5-fold cross-validation to select best-fit Gaussian mixture model parameters according to maximal prediction *R*^2^. For large sample sizes *N* > 10,000, computing a subset of the cross-validation folds is already sufficient to obtain parameter estimates that achieve near-optimal association power, so by default, we compute only enough folds to make predictions on 10,000 test fold samples. This observation holds because absolute prediction *R*^2^ corresponds directly to the power of the corresponding retrospective association test (as measured by increase in *χ*^2^ statistics at truly associated loci, or equivalently, increase in effective sample size; see Online Methods), and the standard error of prediction *R*^2^ estimated via cross-validation scales as the inverse square root of the number of test samples.

#### 1.5 Performance optimization

Beyond the broad algorithmic techniques we have described, the BOLT-LMM software employs a variety of performance optimizations that provide large constant-factor savings (*>*10x) in memory and running time over a straightforward implementation.

##### 1.5.1 Memory

**Computation on raw genotypes.** The key optimization that we implement for memory efficiency is direct computation on raw genotypes. Raw genotypes can be stored in only 2 bits per base, versus 64 bits (8 bytes) for standard double-precision floating point values. However, all previous methods necessarily work with floating point matrices (either the *N × N* GRM or a floating point representation of the *N × M* normalized genotype matrix) because they perform spectral decomposition. In contrast, because BOLT-LMM applies iterative methods that use the genotype matrix only in matrix multiplications, it is enough for us to store the raw genotypes plus lookup tables containing normalization information that we apply on-the-fly. Explicitly, for each SNP, rather than storing a normalized genotype vector in memory, we simply store its raw allele count (0, 1, 2, or missing) for each individual and additionally record its mean allele count and normalization constant. Then, when performing computations involving the SNP, we build a properly normalized SNP vector using the above information. Importantly, this vector can be thrown away after the computation, thus keeping BOLT-LMM’s memory footprint small.

An additional subtlety arises when working with covariates because normalized genotype vectors no longer contain only 4 values after projecting out covariates. In this case, we store the components of each normalized genotype vector along a basis that spans the covariates, and we do the same for the phenotype vector. We treat these “covariate component” values as additional coordinates that we carry along in all computations, so that whenever we need to compute a dot product (the basic computation all of our iterations use), we can do so by taking the usual dot product and subtracting the dot product of the additional covariate component vector.

**Streaming computation of association tests.** In analyses of data sets containing very large numbers of SNPs (e.g., millions of imputed SNPs) that may be stored as real-valued dosages, it is often desirable to compute association statistics for all SNPs but use only a subset of genotyped SNPs in the mixed model. In this situation, we retain memory efficiency by reading and analyzing SNPs *not* used in the mixed model only when performing final computation of retrospective association test statistics. That is, we first compute a set of residual phenotypes *y*_resid-LOCO_ using only the subset of typed SNPs in the model; then, we successively read each test SNP and compute and output its association test, throwing away the SNP after completing the computation. This streaming computation allows us never to store the full set of SNPs in memory.

##### 1.5.2 Computational speed

**Batch computation using optimized matrix subroutines.** The iterative methods we have described reduce the association computation to basic building blocks of vector operations (for variational iteration) and matrix-vector operations (for conjugate gradient iteration). These operations can be performed using optimized implementations of the Basic Linear Algebra Subprograms (BLAS), but all BLAS libraries achieve maximal speed when performing matrix-matrix “Level 3 BLAS” operations. We therefore batch our computations into matrix-matrix multiplications by performing simultaneous updates across SNPs and parameter values.

Conjugate gradient iteration uses Level 2 BLAS operations as written. Variational iteration only uses Level 1 BLAS as written (Section 1.3), so to step up to Level 2 BLAS, we perform block updates of SNP effect sizes. Explicitly, the basic variational iteration consists of re-estimating each SNP effect in turn conditioned on current estimates of other SNP effects, which requires computing dot products of each SNP with the residual phenotype vector. Instead of computing dot products one at a time (a Level 1 BLAS operation), we compute dot products of a block of SNPs at once (a matrix-vector multiplication, which is Level 2 BLAS). When we subsequently update the effect size of each SNP in turn, we just need to be careful to update its dot product to reflect changes that have been made to the residual vector due to previous updates of previous SNPs within the block. We do so using an LD matrix that we precompute for each SNP block.

To step up from Level 2 BLAS to Level 3 BLAS, we make use of the fact that all computationally intensive steps of the BOLT-LMM algorithm require multiple replicates of almost the same computation: Step 1a computes BLUP on different random phenotypes; Step 1b solves a set of similar linear systems for different LOCO reps and calibration SNPs; Step 2a computes variational Bayes assuming different hyperparameter values; Step 2b computes variational Bayes for different LOCO reps. With some care, we can in each case simultaneously perform iterations across the different replicates. The upshot is that by using batch computations, BOLT-LMM performs all O(MN)-time operations using BLAS 3 matrix-matrix multiplications (DGEMM). Our software distribution uses the well-tuned Intel Math Kernel Library (MKL) BLAS implementation.

**Multithreading.** Conveniently, most BLAS implementations also support optimized multithreaded computation on multi-core processors, which are now commonplace. In practice, multithreading rarely decreases computation time by a factor equal to the number of cores used because of various overhead costs, but Level 3 BLAS operations often come close to achieving this theoretically maximal speedup. The BOLT-LMM software supports multithreading through BLAS, and we recommend using this option whenever multiple cores are available. We performed our benchmarking analyses using single-core computation simply to give a fair comparison against methods that do not support multithreading.

**Low-level optimization.** We reduce the overhead of loading raw genotypes and building normalized genotype vectors using a few additional low-level tricks. Instead of looking up normalized genotype values one at a time when loading a SNP, we build an intermediate lookup table that maps all 256 possible values of 4 consecutive genotypes (stored in 1 byte) to a 4-vector of normalized values. When loading these 4-vectors from the lookup table into the normalized SNP vector being built, we use streaming SIMD extensions (SSE instructions) to perform multiple loads at once.

### 2 Theory

#### 2.1 Variational Bayes

In Bayesian analysis, one specifies a probability model over observations and model parameters, often wishing to obtain posterior mean estimates of parameters of interest. Unfortunately, the posterior distribution is typically infeasible to integrate over. The idea of the variational framework (termed “variational Bayes”) is to approximate the posterior distribution with a factorized form that is much easier to work with computationally. The factorized form can be integrated over, allowing calculation of approximate posterior mean estimates. For a full discussion of variational Bayes, we refer to ref. [49]. Previous variational methods for Bayesian linear regression in genetics have been presented in refs. [21,22,35]. Here, we briefly summarize key aspects of variational Bayes for quick reference and to establish notation. We then derive formulas giving the variational iteration used by BOLT-LMM, and we further derive theory establishing equivalence between this family of variational methods and penalized linear regression.

##### 2.1.1 Terminology and notation

We begin by considering a general probability model *p*(*y, β*) over observations *y* and parameters *β*. We assume that *p*(*y, β*) takes into account the prior distributions of *β*, so that *p*(*y, β*) is the joint probability of sampling parameters *β* and observing *y*.

- The true *log likelihood* (LL) of observing *y* is the log of the integral of *p*(*y, β*) over all possible values of the parameters *β*:

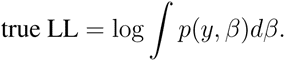 This integral is typically intractable to compute.
• The true *posterior distribution* of the parameters *β* conditional on observing *y* is given by normalizing the joint probability *p*(*y, β*) by the likelihood:

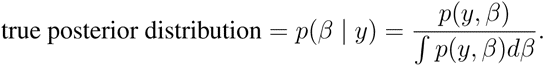
• The *variational approximation to the posterior distribution* is the best approximation of the posterior distribution *p*(*β | y*) with a distribution *q*(*β*) that factors:

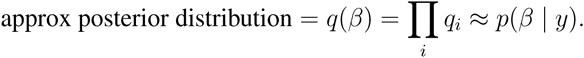 The factors *q_i_* are usually constrained to have simple forms. For instance, if *β* represents a set of individual parameters *β_i_*, each factor *q_i_* may be required to be a function of only one parameter *β_i_*, in which case *q_i_* is an approximate marginal posterior for *β_i_*. This *fully factorized* variational approximation is the approach we consider here.
• The *approximate log likelihood* is a variational lower bound on the true log likelihood, given by:

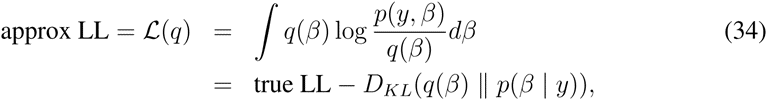

where *D_KL_* denotes the Kullback-Liebler (KL) divergence between probability distributions. Note that the second line says that the gap between the variational lower bound and the true LL is given by the KL divergence between the approximating distribution *q* and the true posterior distribution *p*(*β | y*).

Variational iteration, which we describe in the next section, successively refines the approximation of *p*(*β | y*) with *q*(*β*) by iteratively updating the factors *q_i_*, reducing the KL divergence *D_KL_*(*q*(*β*) || *p*(*β | y*)) and therefore monotonically improving the lower bound *𝓛*(*q*). The point of the iteration is that at convergence, the distribution *q*(*β*) will ideally be a good approximation of the true posterior distribution *p*(*β | y*) that can easily be integrated over (because of its factored form) to perform approximate posterior inference.

Before moving on, we note that it follows from the above that the faithfulness of the approximating distribution *q*(*β*) to the true posterior *p*(*β | y*) determines the accuracy of the resulting approximate posterior inference. Because we only consider approximating distributions *q*(*β*) that factor as 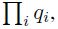 the optimal quality of the approximation depends on the extent to which the posterior distribution *p*(*β | y*) can be approximately factored in the prescribed manner. For Bayesian linear regression, which is our focus here, correlation among regressors (i.e., linkage disequilibrium among markers) tends to pose a challenge for some types of inference using variational methods. For example, when regressors are correlated, the approximate posterior is often too tightly concentrated (i.e., overconfident of parameter localization). However, aggregate inference can still be robust. We refer to ref. [21] for an in-depth exploration of these issues.

##### 2.1.2 General variational iteration

The variational Bayes algorithm iteratively updates each approximate marginal distribution *q_i_* using the update step

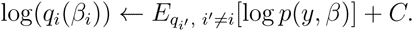

As written, the above update step updates the entire distribution *q_i_*—a *function* of *β_i_*—to its new optimum (as a distribution, optimizing in the sense of calculus of variations). Indeed, the only assumption made by the variational approximation is that the approximate posterior distribution *q*(*β*) factors (i.e., the parameters are assumed to be conditionally independent, conditional on the observed outputs): no assumption is made about the functional form of the factors *q_i_*(*β_i_*). However, the factor distributions *q_i_* are typically characterized by sets of sufficient statistics, so that in each iteration, only the sufficient statistics for factor *i* need to be updated based on the current values of the sufficient statistics for each of the other factors. Moreover, we will show in Section 2.1.3 that in the case of Bayesian linear regression, the factor distributions take the form of conditional posterior distributions. For the Gaussian mixture priors we use in BOLT-LMM, these conditional posteriors retain the Gaussian mixture form, just with different parameters.

Convex optimization theory guarantees the convergence of variational iteration [46]. Moreover, convergence of the approximate log likelihood, which monotonically increases during the iteration, serves as a convergence criterion. At convergence, approximate posterior means of the parameters *β_i_* can be read off from the approximate marginal distributions *q_i_*: for the special case of fully factorized Bayesian linear regression, these values are among the sufficient statistics updated at each iteration.

##### 2.1.3 Variational iteration for Bayesian linear regression

We now specialize to the Bayesian linear regression model

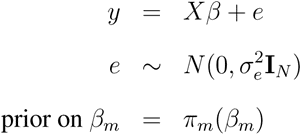

That is, we assume each regression coefficient (i.e., SNP effect) has a prior distribution *π_m_*(*β_m_*) and the response *y* is the sum of regressor effects plus iid Gaussian noise with variance 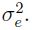 (Note that this noise model does not take into account projecting out covariates as described in Section 1.1; projecting out *C* independent covariates just reduces *N* to *N − C* in the equations below.) The joint probability *p*(*y, β*) of parameters *β* and observations *y* satisfies

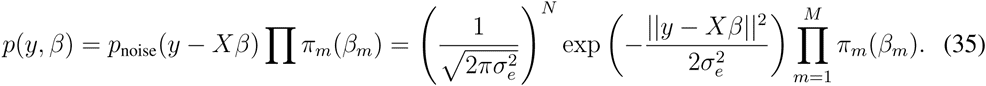

Below we derive the fact that the fully factorized variational approximation matches the iterative conditional update algorithm discussed in Section 1.3. The fully factorized variational approximation takes the form

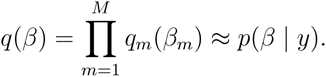

The variational update step for the approximate marginal distribution *q_m_* optimizes the KL divergence *D_KL_*(*q||p*) over all distributions *q_m_*, fixing the other marginals 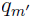 to their current distributions, thereby monotonically increasing the approximate (lower bound) log likelihood *𝓛*(*q*). The update amounts to:

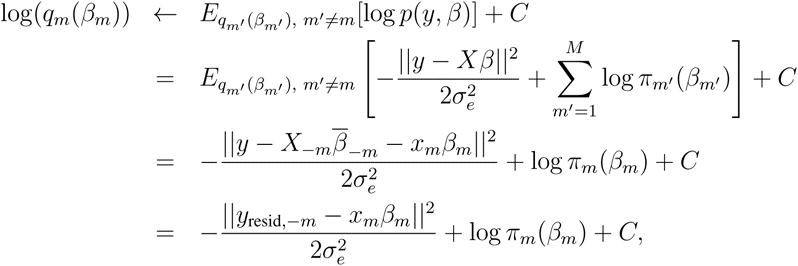

where 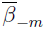 denotes the vector of estimated posterior mean effect sizes at SNPs other than *m* according to the current approximate marginal posterior distributions 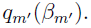 (In the above sequence of equations, we absorb all terms independent of *β_m_* into the constant *C*; the use of “+*C*” in all lines does not imply that the constant is the same from line to line.) To see how the posterior mean estimates 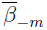 enter the equation in the third line above, consider expanding the quadratic term *||y − Xβ||*^2^ inside the expectation. (Note that taking an expectation over 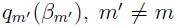 corresponds to integrating over 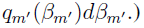

- For terms that are linear in 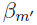 (for 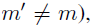), 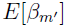 can be replaced with 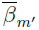 by linearity of expectation over 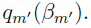
- Terms that are quadratic in 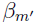 are independent of *β_m_*, so 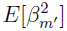 can be replaced by 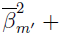 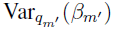 to re-complete the square 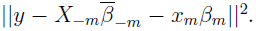 The leftover terms, proportional to 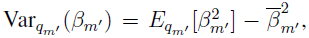 are constant (with respect to *β_m_*) and can be absorbed into the constant term.

Finally, the contributions of the prior terms 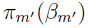 for 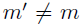 also become additive constants upon taking expectations and can be absorbed into the constant term as well.

Exponentiating the result above gives

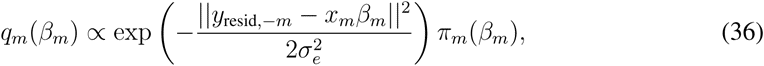

which is simply a conditional posterior distribution for *β_m_* (given the prior *π_m_*(*β_m_*) and conditional on setting all other 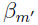 to their variational expected means). That is, for Bayesian linear regression, the optimal approximating marginal distribution *q*(*β_m_*) (making no assumptions about the form of each marginal, only requiring that the approximating posterior fully factorizes) turns out to be the conditional posterior. In particular, as we derive in Section 2.1.4, if the prior has the nice form of a mixture of Gaussians, then the approximating marginal keeps that form (with different means, standard deviations, and weights).

Moreover, the approximate log likelihood *𝓛*(*q*) can be obtained by substituting the joint probability 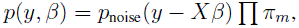 given in equation (35), into equation (34):

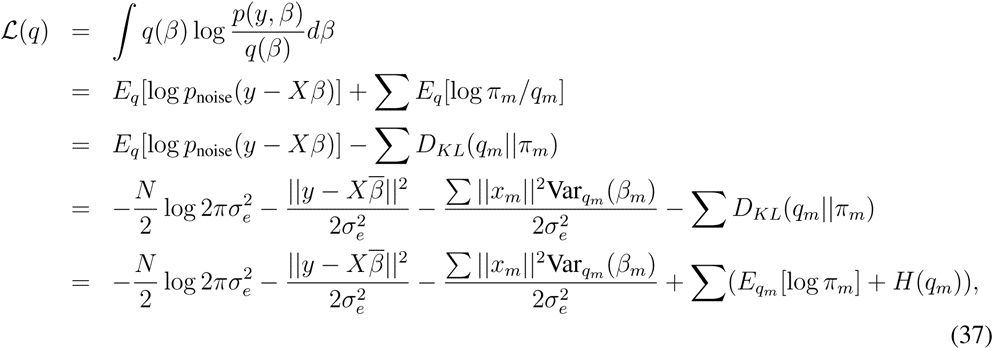

where 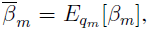 using the same trick of completing the square as above, only now we have to be careful to account for the leftover constant terms involving Var*_q_m__* (*β_m_*). In the last line, *H*(*q_m_*) denotes information entropy. Once again, we note that if *C* independent covariates are projected out from our model as described in Section 1.1, then we simply reduce *N* to *N − C* in equation (37) to account for the lost degrees of freedom.

##### 2.1.4 Update equations for Gaussian mixture prior

We now specialize to Bayesian linear regression with the specific Gaussian mixture prior

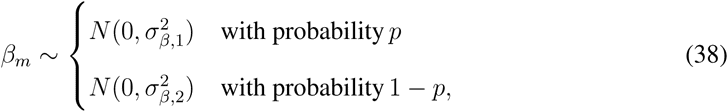

i.e.,

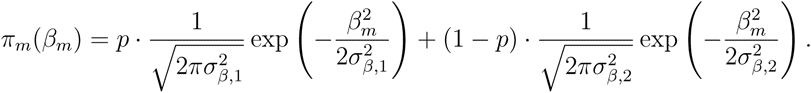

Then the conditional marginal distribution is given by

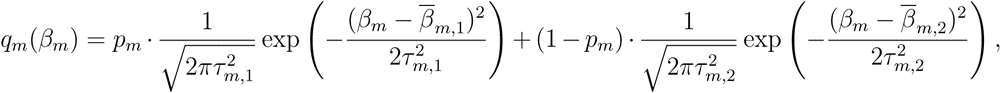

i.e.,

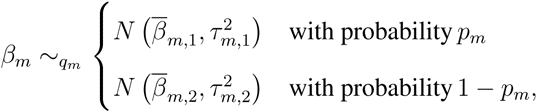

where

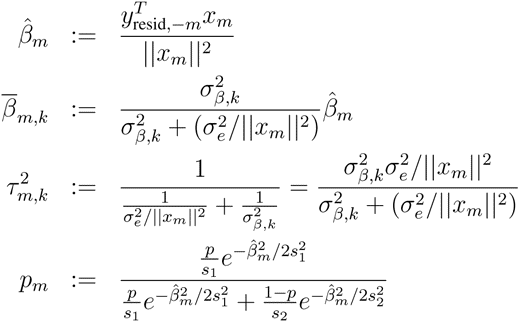

where 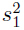 and 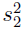 are the variances of the marginal distribution of 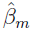 in each case (assuming *y*_resid_*_,−m_* = *x_m_β_m_* + *e* and marginalizing over random *β_m_* and *e*):

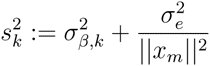

for *k* = 1, 2. These equations explicitly give the update formulas for the BOLT-LMM variational iteration (Step 2 in Section 1.3), in which each estimated SNP effect is set to its conditional posterior mean.

To compute the approximate log likelihood for use in testing convergence of the iteration, it is convenient to consider a reparameterization of the model (following ref. [21]) in which instead of having one parameter *β_m_* for each SNP (with *π_m_*(*β_m_*) a mixture of two Gaussians), we introduce a state parameter *s_m_* ∈ {0, 1} and consider a prior *π_m_*(*β_m_, s_m_*) with:

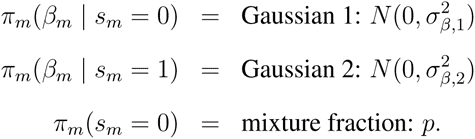

This parameterization gives exactly the same model as the original Bayesian linear regression, with the only difference being that the mixture prior on *β_m_* in equation (38) is replaced by the joint prior distribution *π_m_*(*β_m_, s_m_*) above with the extra hidden state *s_m_*.

We can now apply the variational approach to factorize the approximate posterior distribution over all SNPs—in (*β_m_, s_m_*) pairs—as a product of joint distributions *q_m_*(*β_m_, s_m_*). Each optimal factor distribution *q_m_* has the property that when integrated over *s_m_*, it matches the variational approximation *q_m_*(*β_m_*) from the original parameterization, implying that the estimated posterior means are identical. This parameterization lends itself more easily to computation of KL divergences, however. We have the formula:

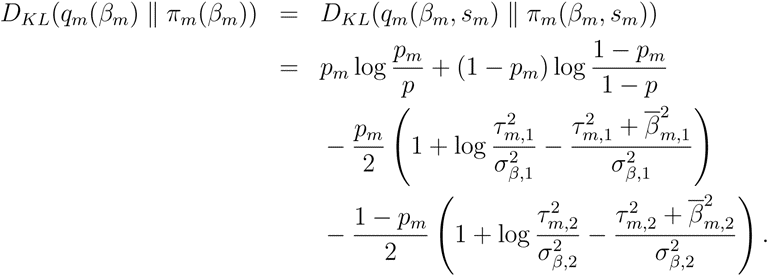

Writing

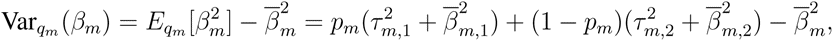

we have all of the terms needed to compute *𝓛*(*q*) according to equation (37).

We note that in the limit *σ_β,_*_2_ *→* 0 of the point-normal prior used by ref. [21], the formulas become

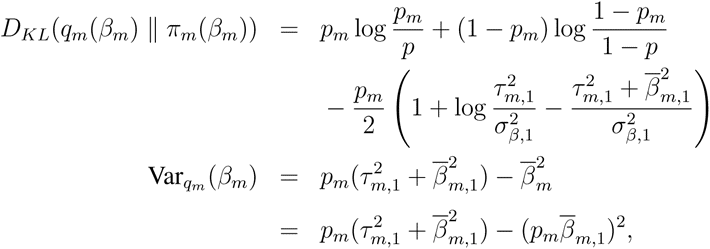

matching ref. [21].

##### 2.1.5 Equivalence with penalized linear regression

We now show that the variational iteration of Section 2.1.3 for Bayesian linear regression is equivalent to coordinate descent applied to a penalized linear regression problem. Equivalently, the fully factorized variational approximation to Bayesian linear regression (for an arbitrary choice of prior on regressor effects) can be recast as applying a transformation to the prior and then finding a posterior mode of the (new) Bayesian linear regression with transformed prior. More precisely, we have the following *equivalence of optimization problems*:

1. Maximize the variational approximate log likelihood (for the fully factored VB approximation to Bayesian linear regression).
2. Minimize the penalized linear regression objective function

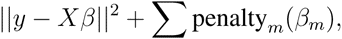

where the penalty is derived from the prior in the original Bayesian linear regression.
3. Find a posterior mode of the Bayesian linear regression with prior

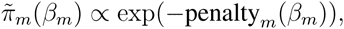

which we can view as a transformation of the original prior *π_m_*(*β_m_*).

We also have the following *equivalence of algorithms*:

1. Variational iteration applied to the original Bayesian linear regression.
2. Coordinate descent applied to the corresponding penalized linear regression.

**Implications for numerical optimization.** The equivalence of variational Bayes with penalized linear regression (in the context of Bayesian linear regression) elucidates some numerical properties of the algorithm that are not immediately apparent. In particular, penalized linear regression is in general a numerically challenging non-convex optimization. We can therefore expect variational iteration to be susceptible to numerical issues such as convergence to local optima. Some methods have tried to address this problem by repeating the iteration for different random choices of update orders or different initialization points [21, 22, 35]. In our simulations, we found that using a single run of the iteration was typically already sufficient for BOLT-LMM to achieve most of the available power gain, however, so we opted to use just a single run to avoid increasing computational cost.

**Mathematical derivation.** As discussed in Section 2.1.1, variational theory gives the following lower bound to the log likelihood, which variational iteration tries to maximize:

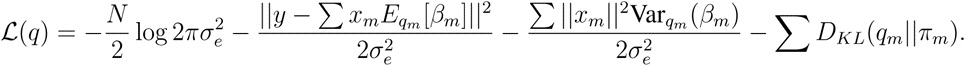

Note that *𝓛*(*q*) is a *functional*, i.e., it evaluates factored distributions *q* = *q*_1_ *· · · q_M_* that attempt to approximate the posterior. We will show that in fact, this maximization is equivalent to optimizing the objective function (over effect size estimates 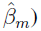 of a penalized linear regression:

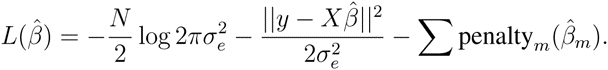

The key is that we can define a mapping 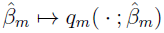 with 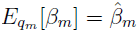 and define

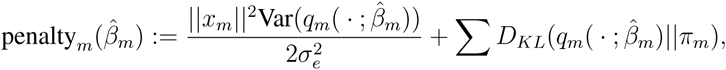

with the property that

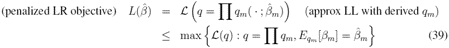

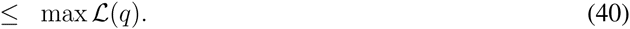

(Note that if the SNPs are normalized such that *||x_m_||* are all equal, then the penalty is independent of *m*.)

The mapping 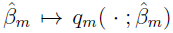 is derived from the variational update step, which we know chooses the *q_m_* that optimizes *𝓛*(*q*) conditional on the current choices of the other marginal distributions. Thus, equality holds in (39) at any convergence “point” of the iteration. Here a convergence “point” really means a choice of approximating distributions *q*_1_,…, *q_M_*; however, if the above mapping exists, these distributions may be parameterized by 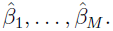 Equality holds in (40) at the global maximizer of the variational approximate log likelihood. It follows that 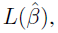 which is upper-bounded by the global maximum of *𝓛*(*q*), attains that global maximum at 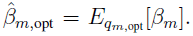 That is, the solution to the penalized linear regression optimization of 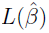 corresponds to the solution to the variational optimization of *𝓛*(*q*): if we could solve the penalized linear regression optimally, we would have the optimal variational Bayes solution.

All that remains is to define the mapping 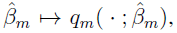 which we obtain indirectly from the variational update step via the key observation that the optimal marginal *q_m_* conditioned on all other marginals depends only on a single statistic: the correlation of *x_m_* with the residual 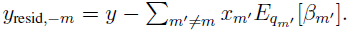 That is, we have:

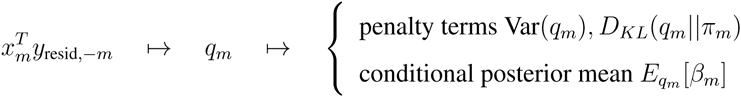

Moreover, the conditional posterior mean *E_q__m_* [*β_m_*] is an increasing function of the correlation statistic 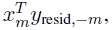 meaning that we can invert the final mapping to obtain 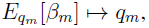 giving the desired mapping 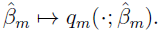 If we wanted to derive an explicit formula for penalty*_m_*(*β_m_*), we could try to invert the mapping 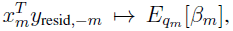 but in general there is probably no closed form.

#### 2.2 Convergence rate of BOLT-LMM iterative computations

Because BOLT-LMM applies iterative methods for numerical linear algebra, its running time depends not only on the cost of matrix operations, which scales linearly with M and N, but also the number of O(MN)-time iterations required for convergence, which is largely determined by the condition number of the phenotypic covariance matrix and in particular increases with sample size (N), heritability, relatedness, and population structure. Simulations show empirically that the number of iterations scales roughly with 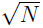 and does not change dramatically within the range of typical values of the other parameters, hence our overall time complexity estimate of *≈*O(MN^1.5^)

Additionally, the high observed-scale heritabilities of case-control ascertained traits and high pseudo-heritabilities of phenotypes in family data, in combination with population structure or relatedness among samples, may reduce the computational efficiency of BOLT-LMM (Supplementary Fig. 2); an avenue for future investigation is to mitigate this effect by including PCs to improve the numerical conditioning of the computation.

### 3 Parameters of mixed model software used in analyses

We ran BOLT-LMM with default options except in analyses investigating power of the Gaussian mixture model, in which we used the `--forceNonInf` option to fit the Gaussian mixture model even in scenarios with no expected power gain.

We ran the 2012-02-10 intel64 release of EMMAX with default options. We ran GEMMA version 0.94 with default options. We ran GCTA version 1.24 with the `--mlma-loco` option for leave-one-chromosome-out analysis. We ran FaST-LMM version 2.07 using all markers to compute the similarity matrix. We ran FaST-LMM-Select version 2.07 with the autoSelect option

`-autoSelectSearchValues`

`“0,1,3,10,30,100,300,1000,3000,10000,30000,100000,300000”`
to test increasing numbers of selected SNPs up to a maximum of all 300,000 SNPs. We benchmarked the precompiled executable distribution of each software package.

**Supplementary Figure 1.**
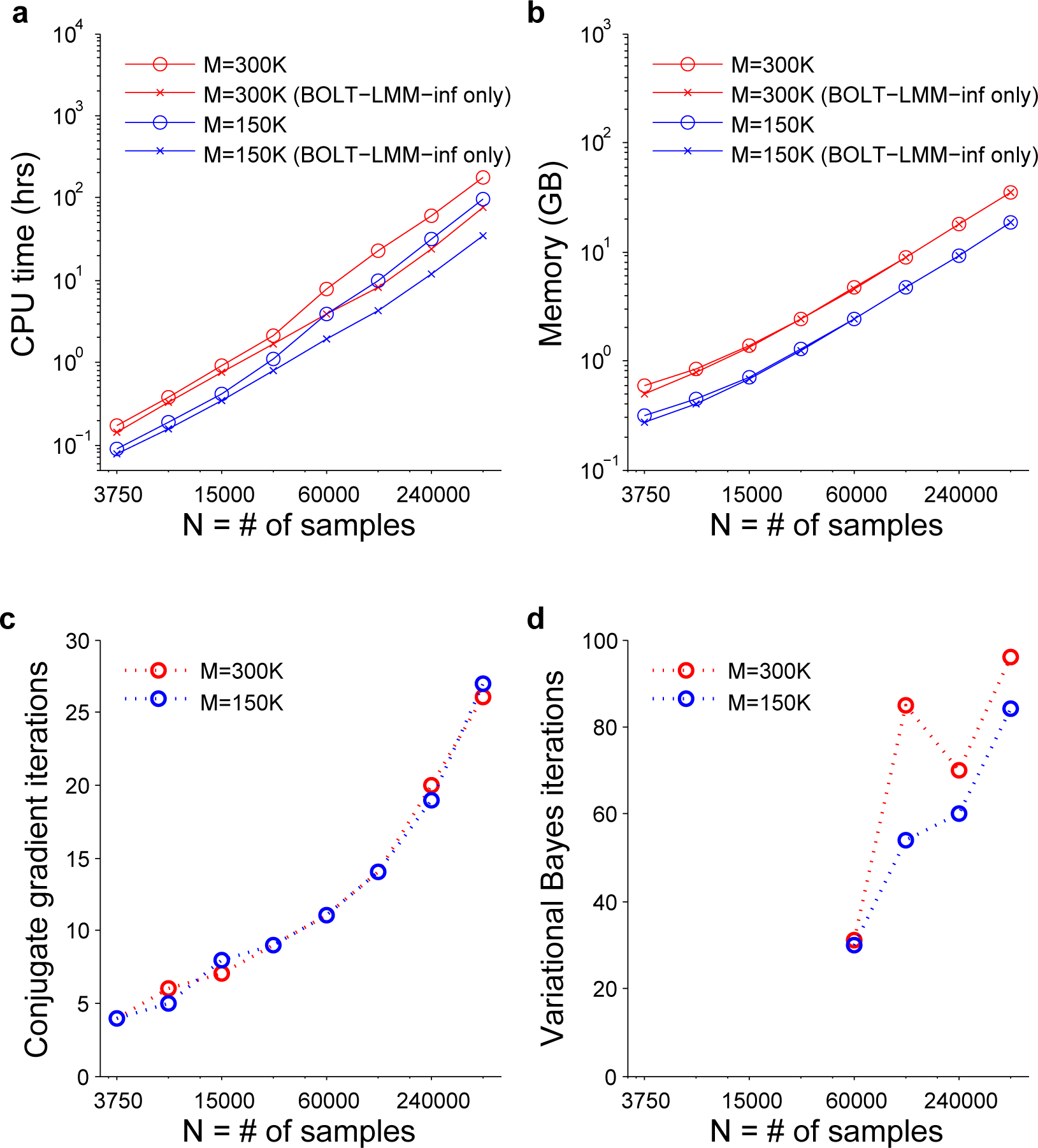
Detailed computational cost metrics from running BOLT-LMM on simulated data sets with increasing sample size (Fig. 1): (**a**) running time, (**b**) memory usage, (**c**) conjugate gradient iterations used in LOCO analysis, (**d**) variational Bayes iterations used in LOCO analysis (max among 22 LOCO reps). Note that the conjugate gradient computation (**c**) is required to compute both the BOLT-LMM-inf and BOLT-LMM statistics, whereas the variational Bayes computation (**d**) is relevant only to the BOLT-LMM statistic. Additionally, BOLT-LMM skips the LOCO variational Bayes computation when estimated improvement in prediction *R*^2^ using the Gaussian mixture model is small (*<*1%); in (**d**), this behavior occurred for N < 60 K. (Note that in such cases, some variational Bayes work is still needed in making this determination; see Online Methods for details.) Each plotted point corresponds to one simulation with M_causal_ = 5000 SNPs explaining *h*^2^_causal_ = 0.2 of phenotypic variance. Reported run times are medians of five identical runs using one core of a 2.27 GHz Intel Xeon L5640 processor.

**Supplementary Figure 2.**
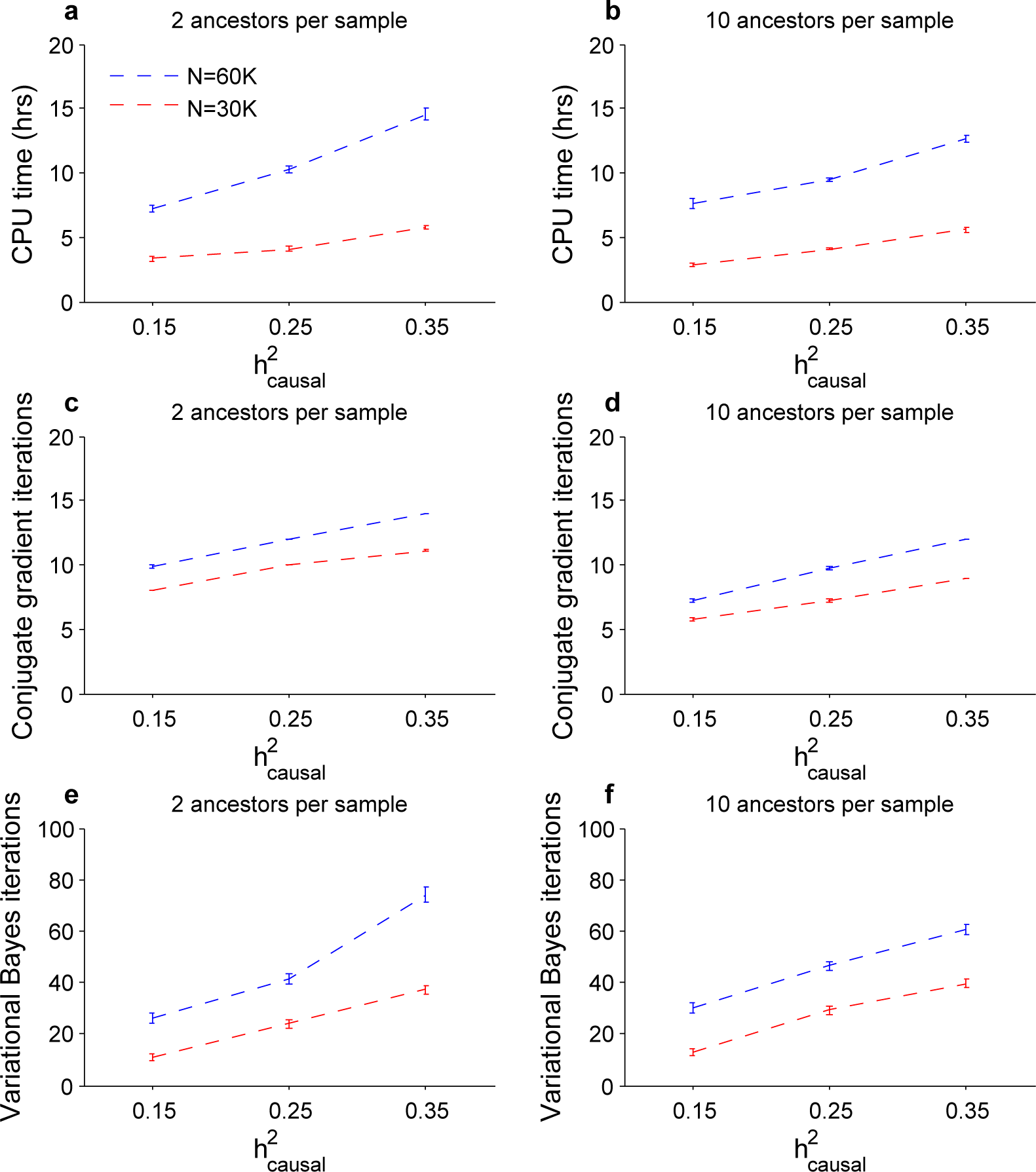
Dependence of BOLT-LMM computational cost metrics (see Supplementary Fig. 1) on heritability explained by genotyped SNPs 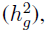 sample size, and sample structure. BOLT-LMM was run on simulated data sets with N = 30,000 or 60,000 samples, each generated as a mosaic of genotype data from 2 (left panels) or 10 (right panels) random “ancestors” from the WTCCC2 data set (N = 15,633, M = 360 K). Phenotypes were simulated with M_causal_ = 5000 SNPs explaining *h*^2^_causal_ = 0.15–0.35 of phenotypic variance. In order to measure variational Bayes iterations (**e**, **f**) used in LOCO analysis for all parameter combinations, BOLT-LMM Gaussian mixture model analysis was run to completion even when estimated improvement in prediction *R*^2^ using the Gaussian mixture model was small (i.e., the default behavior was overriden). Error bars, s.e.m., 10 simulations.

**Supplementary Figure 3.**
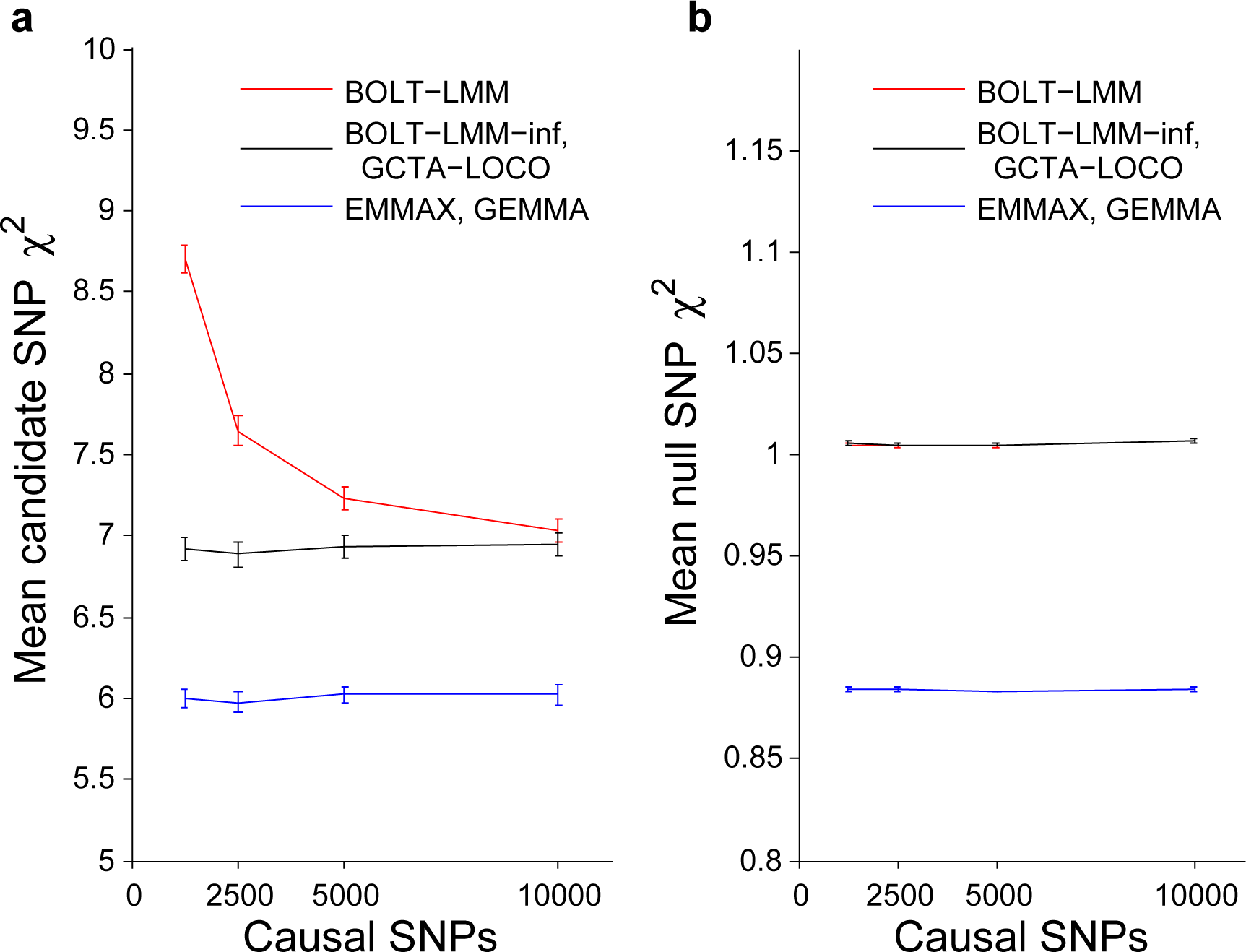
BOLT-LMM increases power to detect associations in simulations while maintaining false positive control. (**a**) Mean *χ*^2^ at causal candidate SNPs. (**b**) Mean *χ*^2^ at null SNPs (i.e., SNPs not in LD with causal SNPs). Simulations used real genotypes from the WTCCC2 data set (N = 15,633, M = 360 K) and simulated phenotypes with the specified number of causal SNPs explaining 50% of phenotypic variance and 60 more candidate SNPs explaining an additional 2% of the variance. Error bars, s.e.m., 100 simulations. Plotted data are for BOLT-LMM, BOLT-LMM-inf, and EMMAX statistics. We verified on the first 5 simulations that the BOLT-LMM-inf and GCTA-LOCO statistics were nearly identical and that the EMMAX and GEMMA statistics were nearly identical (Supplementary Table 6).

**Supplementary Figure 4.**
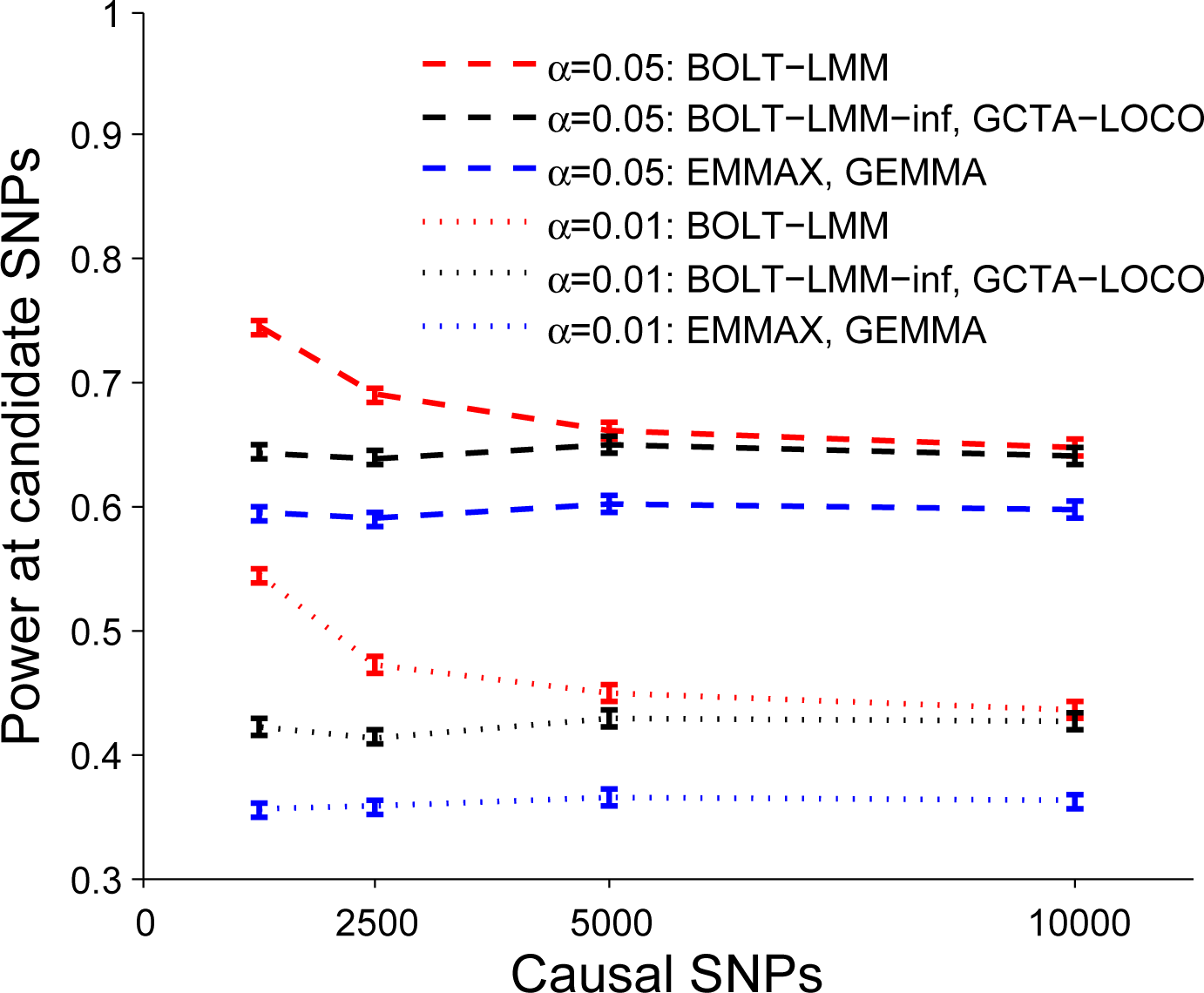
Power of BOLT-LMM and existing mixed model association methods to detect causal candidate SNPs in the simulations of Fig. 3 (real genotypes from the WTCCC2 data set with N = 15,633 and M = 360 K, simulated phenotypes with varying numbers of causal SNPs). Error bars, s.e.m., 100 simulations.

**Supplementary Figure 5.**
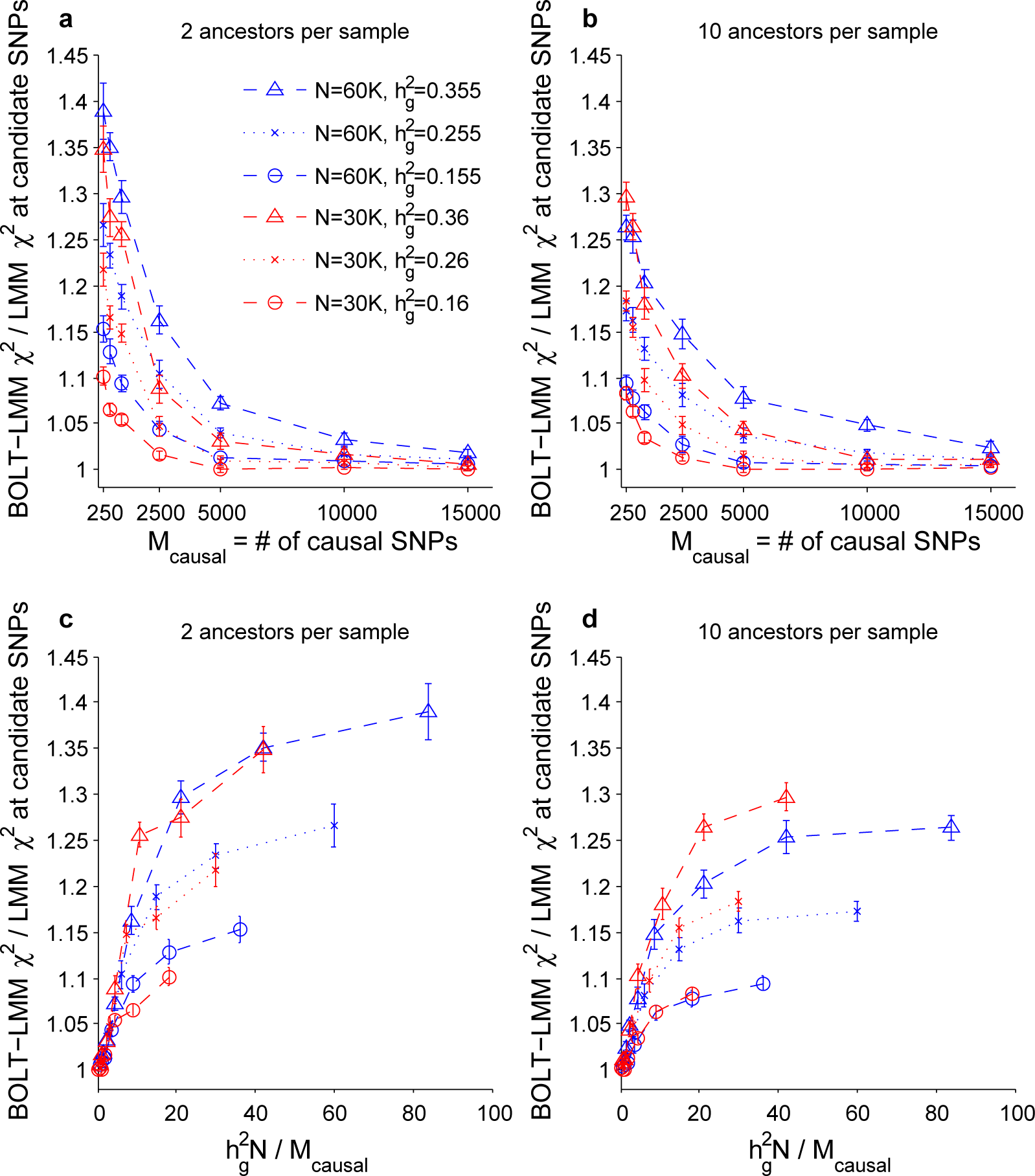
Power increase of BOLT-LMM Gaussian mixture model association analysis over standard mixed model analysis as a function of sample size, number of causal SNPs, and heritability explained by genotyped SNPs 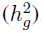. Plotted are ratios of mean *χ*^2^ BOLT-LMM vs. BOLT-LMM-inf statistics at causal candidate SNPs as a function of (**a**) M_causal_ or (**b**) 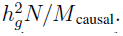 This metric is equivalent to increase in effective sample size using the Gaussian mixture model. Data sets used N = 30,000 or 60,000 simulated individuals, each generated as a mosaic of genotype data from 2 (left panels) or 10 (right panels) random “ancestors” from the WTCCC2 data set (N = 15,633, M = 360 K). Phenotypes were simulated with M_causal_ = 2500–15,000 SNPs explaining *h*^2^_causal_ = 0.15–0.35 of phenotypic variance and 60 more candidate SNPs explaining an additional 0.005 (N = 60 K) or 0.01 (N = 30 K) of the variance. In all simulations, both BOLT-LMM and BOLT-LMM-inf statistics were properly calibrated (mean *χ*^2^ = 1.00–1.01 at null SNPs). Error bars, s.e.m., 10 simulations.

**Supplementary Figure 6.**
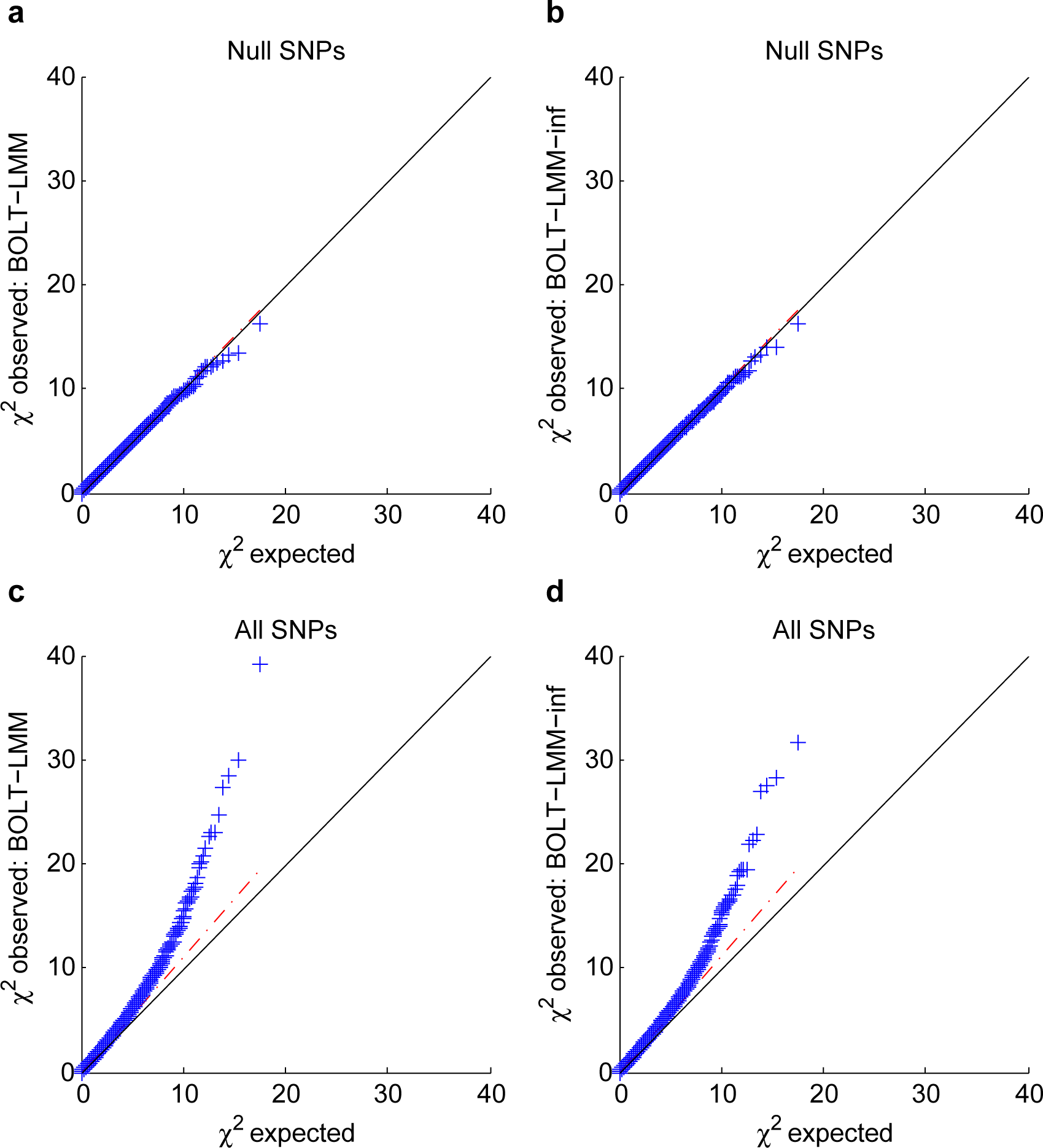
Q-Q plots of BOLT-LMM *χ*^2^ statistics (**a**, **c**) and BOLT-LMM-inf *χ*^2^ statistics (**b**, **d**). The observed quantiles of both association statistics at null SNPs (**a**, **b**) match theoretical *χ*^2^ quantiles. The observed test statistics over all SNPs (**c**, **d**) show lift-off consistent with polygenicity as simulated. Data shown are from one simulation using the setup of Fig. 3 (real genotypes from the WTCCC2 data set with N = 15,633 and M = 360 K, simulated phenotypes with 5060 causal SNPs explaining 0.52 of phenotypic variance). To reduce clutter, 5% of the SNPs are plotted.

**Supplementary Figure 7.**
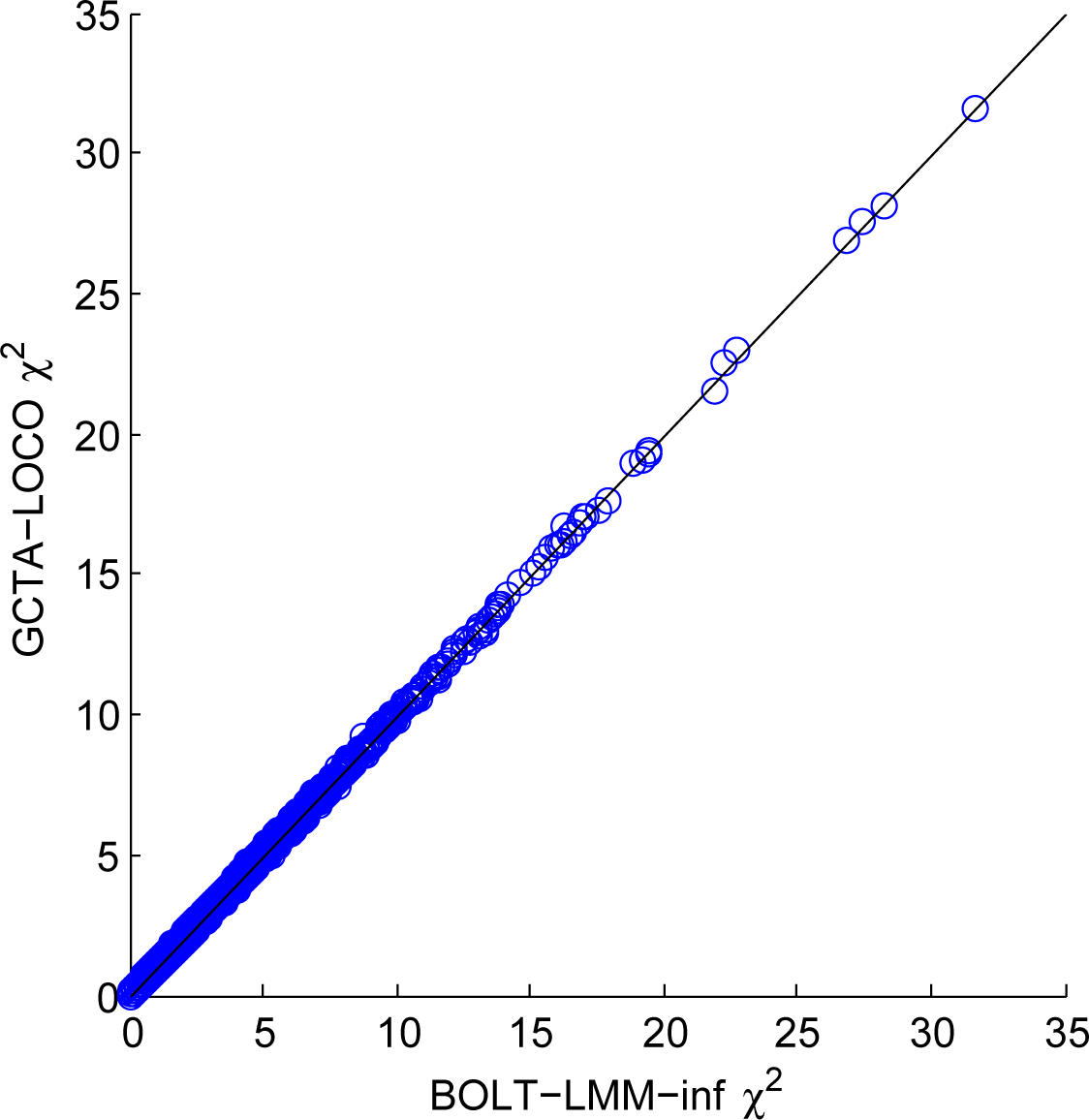
Scatter plot of BOLT-LMM-inf vs. GCTA-LOCO *χ*^2^ statistics. BOLT-LMM-inf and GCTA-LOCO are expected to differ slightly because GCTA-LOCO is the standard prospective statistic whereas BOLT-LMM-inf is a retrospective statistic. Data shown are from one simulation using the setup of Fig. 3 (real genotypes from the WTCCC2 data set, simulated phenotypes with 5060 causal SNPs explaining 0.52 of phenotypic variance). To reduce clutter, 5% of the SNPs are plotted.

**Supplementary Table 1.** Computational performance of mixed model association methods

| N | BOLT-LMM | BOLT-LMM-inf | GCTA-LOCO | EMMAX | GEMMA | FaST-LMM | FaST-LMM-Select |
| --- | --- | --- | --- | --- | --- | --- | --- |
| 3,750 | 0.171 hr / 0.582 GB | 0.141 hr / 0.5 GB | 1.15 hr / 6.31 GB | 1.78 hr / 3.77 GB | 3.28 hr / 0.323 GB | 1.22 hr / 25.4 GB | 4.24 hr / 25.5 GB |
| 7,500 | 0.385 hr / 0.847 GB | 0.324 hr / 0.771 GB | 5.58 hr / 15.8 GB | 8.79 hr / 3.78 GB | 12.2 hr / 0.953 GB | 3.34 hr / 50.9 GB | 15.8 hr / 54.9 GB |
| 15,000 | 0.91 hr / 1.39 GB | 0.78 hr / 1.32 GB | 31.4 hr / 44.8 GB | 32.1 hr / 13.4 GB | 50 hr / 3.47 GB | NA<sup>a</sup> | NA<sup>a</sup> |
| 30,000 | 2.11 hr / 2.45 GB | 1.67 hr / 2.41 GB | NA<sup>a</sup> | 129 hr / 53.7 GB | NA<sup>b</sup> | NA<sup>a</sup> | NA<sup>a</sup> |
| 60,000 | 7.93 hr / 4.62 GB | 3.94 hr / 4.59 GB | NA<sup>a</sup> | NA<sup>a</sup> | NA<sup>b</sup> | NA<sup>a</sup> | NA<sup>a</sup> |
| 120,000 | 23.4 hr / 8.96 GB | 8.07 hr / 8.96 GB | NA<sup>a</sup> | NA<sup>a</sup> | NA<sup>b</sup> | NA<sup>a</sup> | NA<sup>a</sup> |
| 240,000 | 60.8 hr / 17.7 GB | 24.1 hr / 17.7 GB | NA<sup>a</sup> | NA<sup>a</sup> | NA<sup>b</sup> | NA<sup>a</sup> | NA<sup>a</sup> |
| 480,000 | 177 hr / 34.3 GB | 76 hr / 35.1 GB | NA<sup>a</sup> | NA<sup>a</sup> | NA<sup>b</sup> | NA<sup>a</sup> | NA<sup>a</sup> |

**Supplementary Table 2.** BOLT-LMM increases power to detect associations in simulations

| (a) | $M_{\text{causal}}$ | PCA | BOLT-LMM-inf | BOLT-LMM |
| --- | --- | --- | --- | --- |
|  | 1250 | 6.55 (0.07) | 6.92 (0.07) | 8.71 (0.08) |
|  | 2500 | 6.52 (0.07) | 6.89 (0.08) | 7.65 (0.09) |
|  | 5000 | 6.56 (0.07) | 6.94 (0.07) | 7.23 (0.07) |
|  | 10000 | 6.56 (0.07) | 6.94 (0.07) | 7.04 (0.07) |

| (b) | $h^2_{\text{causal}}$ | PCA | BOLT-LMM-inf | BOLT-LMM |
| --- | --- | --- | --- | --- |
|  | 0.3 | 6.41 (0.07) | 6.54 (0.07) | 6.71 (0.07) |
|  | 0.5 | 6.52 (0.07) | 6.89 (0.08) | 7.65 (0.09) |
|  | 0.7 | 6.63 (0.08) | 7.39 (0.09) | 9.71 (0.11) |

| (c) | $N$ | PCA | BOLT-LMM-inf | BOLT-LMM |
| --- | --- | --- | --- | --- |
|  | 5211 | 2.83 (0.04) | 2.88 (0.04) | 2.90 (0.04) |
|  | 10422 | 4.71 (0.06) | 4.90 (0.06) | 5.14 (0.06) |
|  | 15633 | 6.52 (0.07) | 6.89 (0.08) | 7.65 (0.09) |

These tables provide the numerical data (mean *χ*^2^ statistics at candidate causal SNPs) plotted in Fig. 2.

**Supplementary Table 3.** Calibration of BOLT-LMM statistics in simulations with varying genetic architectures and sample sizes

| | $M_{\text{causal}}$ | PCA | BOLT-LMM-inf | BOLT-LMM |
| --- | --- | --- | --- | --- |
| (a) | 1250 | 1.009 (0.001) | 1.006 (0.001) | 1.005 (0.001) |
|  | 2500 | 1.008 (0.001) | 1.005 (0.001) | 1.004 (0.001) |
|  | 5000 | 1.007 (0.001) | 1.005 (0.001) | 1.004 (0.001) |
|  | 10000 | 1.009 (0.001) | 1.007 (0.001) | 1.007 (0.001) |

| | $h^2_{\text{causal}}$ | PCA | BOLT-LMM-inf | BOLT-LMM |
| --- | --- | --- | --- | --- |
| (b) | 0.3 | 1.005 (0.001) | 1.004 (0.001) | 1.005 (0.001) |
|  | 0.5 | 1.008 (0.001) | 1.005 (0.001) | 1.004 (0.001) |
|  | 0.7 | 1.011 (0.001) | 1.009 (0.001) | 1.000 (0.001) |

| | $N$ | PCA | BOLT-LMM-inf | BOLT-LMM |
| --- | --- | --- | --- | --- |
| (c) | 5211 | 1.002 (0.001) | 1.004 (0.001) | 1.004 (0.001) |
|  | 10422 | 1.005 (0.001) | 1.005 (0.001) | 1.005 (0.001) |
|  | 15633 | 1.008 (0.001) | 1.005 (0.001) | 1.004 (0.001) |

We report mean *χ*^2^ statistics at null SNPs in the simulations of Fig. 2. The very slight inflation of BOLT-LMM and BOLT-LMM-inf arises from the use of approximate variance parameter estimates and from the fact that standard mixed model methods eliminate the effects of stratification nearly perfectly but not completely (Supplementary Table 4).

**Supplementary Table 4.**
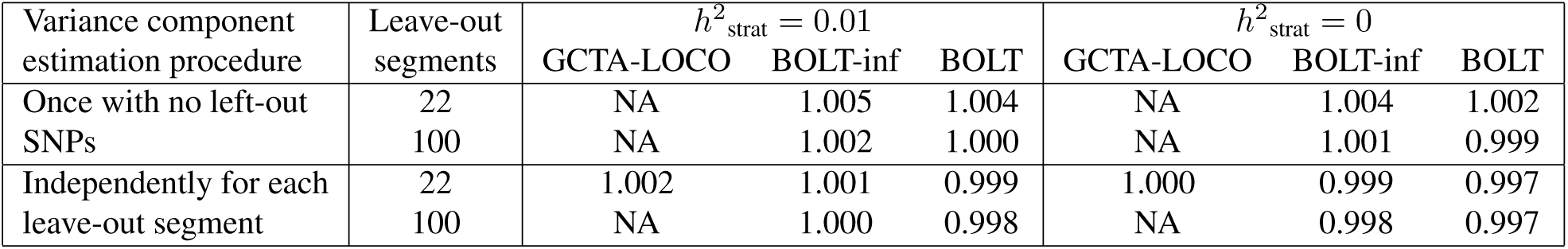
Calibration of BOLT-LMM statistics using different variance component estimation procedures

Mean GCTA-LOCO, BOLT-LMM-inf, and BOLT-LMM chi-squared statistics at null SNPs computed using different methods for variance component analysis and different numbers of leave-out segments. We used the same simulation setup as in Fig. 2 (real genotypes from the WTCCC2 data set, simulated phenotypes with 2560 causal SNPs explaining 0.52 of phenotypic variance). We performed two sets of simulations: one with environmental stratification *h*^2^_strat_ explaining 0.01 of the variance, and the other with no environmental stratification. Thus, the top left entries of 1.005 for BOLT-LMM-inf and 1.004 for BOLT-LMM correspond to the calibration results in Supplementary Table 3 for *M* _causal_ = 2500, *h*^2^_causal_ = 0.5, *N* = 15,633 (appearing in each of Supplementary Table 3a,b,c). Data shown are from 100 simulations, which gave a standard error of 0.001 for all numbers above. By default, the BOLT-LMM algorithm estimates variance parameters once using all SNPs and then reuses the variance estimates when computing association statistics, leaving each of the 22 chromosomes out in turn. This procedure is an approximation of the theoretically precise method of re-estimating variance parameters once per leave-out segment, which the BOLT-LMM software offers as an alternative option (Online Methods). The quality of the approximation can also be improved by subdividing chromosomes for leave-out analysis; we compare results with 22 vs. 100 leave-out segments, which the BOLT-LMM software allows as well (Online Methods). GCTA-LOCO re-estimates variance components once for each of 22 left-out chromosomes. These simulations demonstrate that very slight (0.1–0.5%) inflation of *χ*^2^ test statistics can stem from two causes: (1) reusing the same variance parameter estimates across all LOCO reps rather than re-estimating them for each LOCO rep; and (2) near-complete but imperfect correction for stratification by mixed model methods [50]. The first source of slight inflation is specific to the BOLT-LMM approximation procedure but can be reduced by partitioning the genome into finer leave-out segments or eliminated by refitting variance parameters for each LOCO rep at a small increase in computational cost (Online Methods). This inflation does not increase with sample size (Supplementary Table 3c). The second source of slight inflation is shared by all mixed model methods, as evidenced by the 0.2% inflation of GCTA-LOCO in the *h*^2^_strat_ = 0.01 simulation, and scales with sample size and proportion of variance explained by ancestry in the same manner as inflation caused by genuine polygenic effects (see Supplementary Table 2 of ref. [12]). To completely eliminate this source of very slight inflation, it is necessary to either include principal components as fixed effects or include a second variance component that models ancestry [12, 50–52]. However, we believe that this very slight inflation is not a significant problem for standard mixed model methods or BOLT-LMM.

**Supplementary Table 5.**
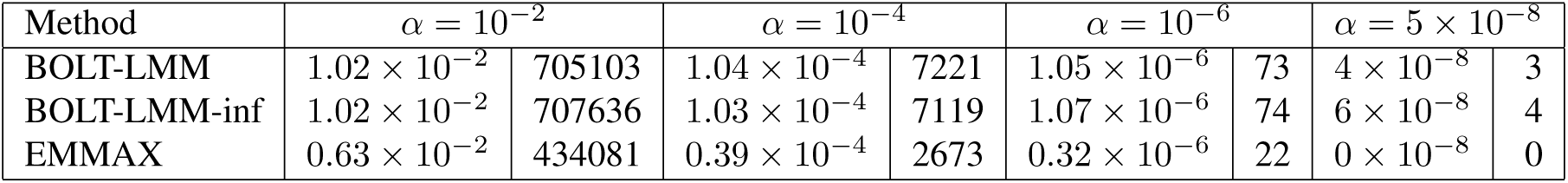
Type I error of BOLT-LMM and EMMAX association tests in simulations

| Method | $\alpha = 10^{-2}$ | | $\alpha = 10^{-4}$ | | $\alpha = 10^{-6}$ | | $\alpha = 5 \times 10^{-8}$ | |
| --- | --- | --- | --- | --- | --- | --- | --- | --- |
| BOLT-LMM | $1.02 \times 10^{-2}$ | 705103 | $1.04 \times 10^{-4}$ | 7221 | $1.05 \times 10^{-6}$ | 73 | $4 \times 10^{-8}$ | 3 |
| BOLT-LMM-inf | $1.02 \times 10^{-2}$ | 707636 | $1.03 \times 10^{-4}$ | 7119 | $1.07 \times 10^{-6}$ | 74 | $6 \times 10^{-8}$ | 4 |
| EMMAX | $0.63 \times 10^{-2}$ | 434081 | $0.39 \times 10^{-4}$ | 2673 | $0.32 \times 10^{-6}$ | 22 | $0 \times 10^{-8}$ | 0 |

Type I error rates and counts for test statistics at null SNPs in the simulations of Fig. 3 (real genotypes from the WTCCC2 data set, simulated phenotypes with causal SNPs explaining 52% of phenotypic variance and environmental stratification explaining 1% of phenotypic variance). To increase statistical resolution, we combined data from all 400 simulations (with 1310, 2560, 5060, and 10060 causal SNPs, 100 simulations each) for a total of 69,293,600 hypothesis tests. Type I error was statistically indistinguishable between different subsets of 100 simulations. The very slight upward bias of actual vs. expected Type I error rates for the BOLT-LMM-inf and BOLT-LMM association tests is explained by very slight (0.1–0.5%) inflation of *χ*^2^ test statistics (Supplementary Table 4), which can be mitigated by performing a more precise variance parameter calculation if desired (Online Methods).

**Supplementary Table 6.** Correlations between mixed model statistics computed by various mixed model methods under the infinitesimal model

| $R^2$ (s.e.m.) | GCTA-LOCO | GEMMA |
| --- | --- | --- |
| BOLT-LMM-inf | 0.999471 (0.000017) | 0.962928 (0.000916) |
| EMMAX | 0.963326 (0.000969) | 0.999997 (0.000000) |

Squared correlation coefficients between *χ*^2^ association statistics over all SNPs computed by various mixed model methods using the simulation setup of Fig. 3 (real genotypes from the WTCCC2 data set, simulated phenotypes with 5060 causal SNPs explaining 0.52 of phenotypic variance). Means and standard errors over 5 simulations are reported. BOLT-LMM-inf and GCTA-LOCO statistics both avoid proximal contamination via LOCO analysis but are expected to differ slightly because GCTA-LOCO is the standard prospective statistic whereas BOLT-LMM-inf is a retrospective statistic. EMMAX and GEMMA are both susceptible to proximal contamination but are expected to differ slightly because GEMMA is an exact statistic whereas EMMAX is an approximate statistic.

**Supplementary Table 7.** Comparison of *χ*^2^ statistics computed by various methods at known SNPs for WGHS phenotypes

| Phenotype | # known SNPs | # tagged SNPs | # $p < 0.05$ PCA & LMM | Mean log fold $\chi^2$ LMM / PCA | # $p < 0.05$ PCA & BOLT | Mean log fold $\chi^2$ BOLT / PCA |
| --- | --- | --- | --- | --- | --- | --- |
| ApoB | 59 (ref. [53]) | 57 | 40 | 2.3 (1.4) | 40 | 9.9 (2.1) |
| LDL | 59 (ref. [53]) | 57 | 45 | 1.4 (1.1) | 45 | 8.7 (1.7) |
| HDL | 72 (ref. [53]) | 72 | 55 | 3.4 (1.2) | 55 | 9.4 (1.8) |
| Cholesterol | 73 (ref. [53]) | 71 | 53 | 1.8 (1.1) | 54 | 5.5 (1.2) |
| Triglycerides | 41 (ref. [53]) | 39 | 29 | -0.6 (1.8) | 29 | 1.7 (2.0) |
| Height | 180 (ref. [54]) | 170 | 131 | 6.0 (1.5) | 134 | 6.2 (1.6) |
| BMI | 32 (ref. [55]) | 31 | 18 | 4.2 (1.8) | 18 | 3.8 (1.8) |
| SystolicBP | 20 (ref. [56]) | 20 | 11 | 2.6 (1.9) | 11 | 2.6 (1.9) |
| DiastolicBP | 19 (ref. [56]) | 19 | 9 | 3.2 (1.8) | 9 | 3.0 (2.0) |

The first two data columns provide additional information about known SNPs used in the power comparison of Supplementary Table 8: # known SNPs, number of genome-wide significant associated SNPs reported in largest GWAS to date; # tagged SNPs, number of such SNPs with an *R*^2^ ≥ 0.2 tagging SNP typed in WGHS. (Note that for the ApoB phenotype, we used known SNPs for the closely related LDL phenotype.) Sums of *χ*^2^ statistics compared in Supplementary Table 8 were computed across the WGHS-typed tagging SNPs.

We also report here one additional metric for increase of power: Mean log fold-change in *χ*^2^ statistics at tagging SNPs. This metric weights all tagging SNPs evenly (Supplementary Table 8), whereas comparing sums of *χ*^2^ statistics weights stronger associations more heavily. Because log fold-change is sensitive to noise from non-replicating SNPs, we restrict to tagging SNPs with at least nominal significance (*p* < 0.05) according to both methods being compared. Methods: PCA, linear regression using 10 principal components as covariates; LMM, BOLT-LMM-inf; BOLT, BOLT-LMM. Errors, s.e.m.

**Supplementary Table 8.** BOLT-LMM increases power to detect associations for WGHS phenotypes

| Phenotype | $h^2_{\text{pseudo}}$ | Prediction $R^2$ | | | | $\chi^2$ incr. at known loci | |
| --- | --- | --- | --- | --- | --- | --- | --- |
| | | PCA (%) | LMM (%)<br>Actual, Expected | BOLT (%) | $p$-value<br>BOLT>PCA | LMM vs.<br>PCA (%) | BOLT vs.<br>PCA (%) |
| ApoB | 0.24 | 0.0 | 1.5 (0.1), 1.7 | 10.4 (0.3) | 4.3E-6 | 1.5 (0.6) | 10.0 (1.2) |
| LDL | 0.18 | 0.1 | 1.0 (0.1), 1.0 | 7.5 (0.2) | 2.8E-6 | 1.4 (0.6) | 8.1 (0.8) |
| HDL | 0.22 | 0.1 | 1.7 (0.1), 1.5 | 6.7 (0.2) | 2.3E-5 | 1.7 (1.5) | 7.1 (2.2) |
| Cholesterol | 0.20 | 0.1 | 1.1 (0.1), 1.1 | 5.9 (0.1) | 1.3E-6 | 1.7 (0.7) | 6.9 (0.8) |
| Triglycerides | 0.18 | 0.0 | 1.2 (0.1), 1.0 | 4.4 (0.3) | 0.00014 | 0.4 (1.0) | 3.1 (1.2) |
| Height | 0.47 | 1.9 | 7.7 (0.4), 6.4 | 9.5 (0.3) | 1.9E-6 | 6.4 (1.3) | 7.6 (1.4) |
| BMI | 0.23 | 0.0 | 1.6 (0.1), 1.5 | 2.1 (0.1) | 8.5E-5 | 1.3 (1.4) | 1.3 (1.5) |
| SystolicBP | 0.17 | 0.2 | 1.0 (0.1), 0.8 | 1.0 (0.1) | 0.00014 | 3.6 (2.2) | 3.6 (2.2) |
| DiastolicBP | 0.14 | 0.0 | 0.7 (0.0), 0.5 | 0.7 (0.0) | 0.00017 | 2.9 (1.7) | 2.7 (1.6) |

We report relative power of different association tests using two roughly equivalent metrics: comparison of *χ*^2^ statistics at known loci, a direct but noisy approach, and out-of-sample prediction *R*^2^ based on the underlying model (both plotted in Figure 3. For reference, we also report pseudo-heritability (*h*^2^_pseudo_), as estimated by BOLT-LMM, and expected prediction *R*^2^ for LMM [31]. Methods: PCA, linear regression using 10 principal components; LMM, standard (infinitesimal) mixed model; BOLT, BOLT-LMM Gaussian mixture model. Actual prediction *R*^2^ values are from 5-fold cross-validation: predictions for each left-out fold were computed by fitting all SNP effects simultaneously (for mixed model methods) or estimating covariate effects (for PCA) using the training folds. Note in particular that for PCA, only covariate effects, not SNP effects, were used for prediction. Standard errors are across folds. Expected prediction *R*^2^ for LMM was computed using N = 23, 294 × 4*/*5 (taking into account the 5-fold cross-validation), M_eff_ = 60,000, and 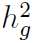 = *h*^2^_pseudo_ estimated by BOLT-LMM, given that the WGHS data set contains little relatedness. (Note that for height, actual prediction *R*^2^ for LMM is slightly higher than expected because PCs explain a non-negligible amount of variance.) Significance for BOLT-LMM > PCA prediction *R*^2^ was assessed using a one-sided paired t-test across folds. Percent increases in *χ*^2^ statistics computed by various methods across known loci are comparisons between sums of *χ*^2^ statistics over typed SNPs in highest LD with published associated SNPs from the largest GWAS to date (Supplementary Table 7). Standard errors are jackknife estimates.

**Supplementary Table 9.** Effect size mixture parameters chosen by BOLT-LMM for WGHS phenotypes

| Phenotype | $h^2_{\text{pseudo}}$ | $f_2$ | $p$ |
| --- | --- | --- | --- |
| ApoB | 0.24 | 0.1 | 0.01 |
| LDL | 0.18 | 0.1 | 0.01 |
| HDL | 0.22 | 0.1 | 0.01 |
| Cholesterol | 0.20 | 0.1 | 0.01 |
| Triglycerides | 0.18 | 0.1 | 0.01 |
| Height | 0.47 | 0.3 | 0.02 |
| BMI | 0.23 | 0.5 | 0.01 |
| SystolicBP | 0.17 | 0.5 | 0.5 |
| DiastolicBP | 0.14 | 0.5 | 0.01 |

We report the best-fit mixture-of-Gaussians prior on SNP effect sizes determined by BOLT-LMM using cross-validation to optimize out-of-sample prediction *R*^2^. The spike and slab mixture of Gaussians is parameterized by the total variance attributed to the combined Gaussian mixture along with two mixture parameters: *f*_2_, the proportion of variance allotted to the spike component (small-effect SNPs), and *p*, the probability that a SNP effect is drawn from the slab component (large-effect SNPs). Note that *f*_2_ = 0.5*, p* = 0.5 corresponds to the infinitesimal model: when *f*_2_ = 1 *-p*, the two Gaussians are identical and the mixture is degenerate.

**Supplementary Table 10.**
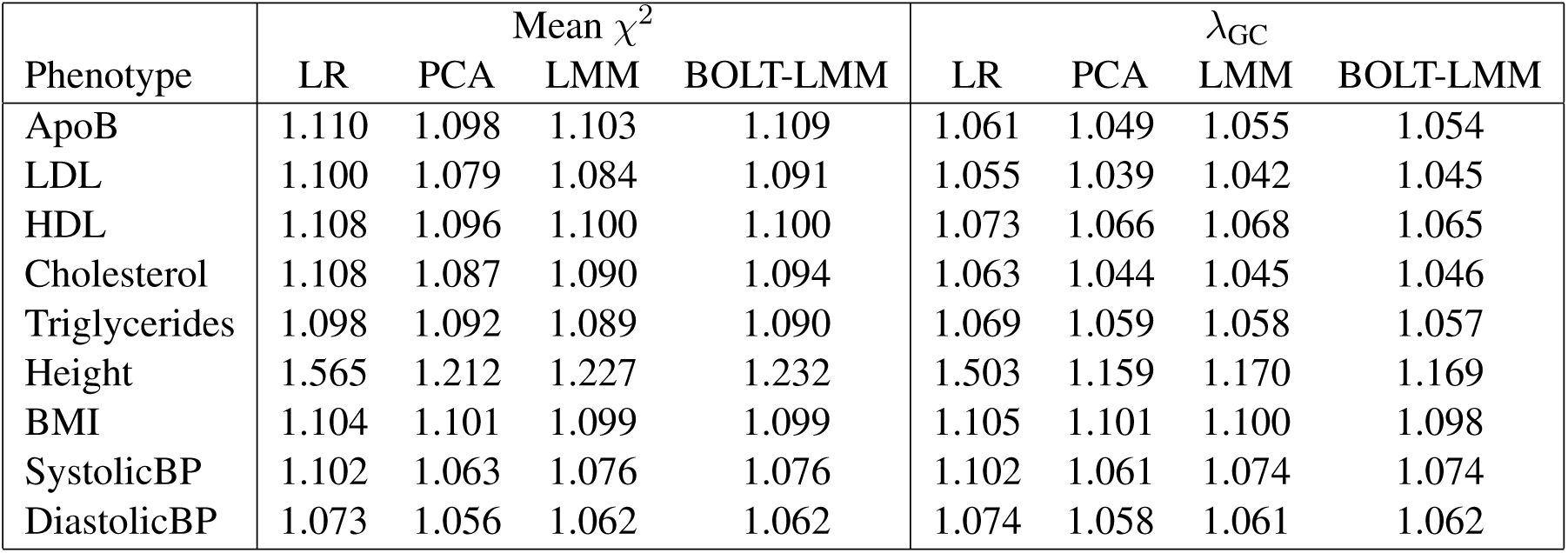
Calibration of *χ*^2^ statistics computed by various methods for WGHS phenotypes

| Phenotype | Mean $\chi^2$ | | | | $\lambda_{GC}$ | | | |
| --- | --- | --- | --- | --- | --- | --- | --- | --- |
|  | LR | PCA | LMM | BOLT-LMM | LR | PCA | LMM | BOLT-LMM |
| ApoB | 1.110 | 1.098 | 1.103 | 1.109 | 1.061 | 1.049 | 1.055 | 1.054 |
| LDL | 1.100 | 1.079 | 1.084 | 1.091 | 1.055 | 1.039 | 1.042 | 1.045 |
| HDL | 1.108 | 1.096 | 1.100 | 1.100 | 1.073 | 1.066 | 1.068 | 1.065 |
| Cholesterol | 1.108 | 1.087 | 1.090 | 1.094 | 1.063 | 1.044 | 1.045 | 1.046 |
| Triglycerides | 1.098 | 1.092 | 1.089 | 1.090 | 1.069 | 1.059 | 1.058 | 1.057 |
| Height | 1.565 | 1.212 | 1.227 | 1.232 | 1.503 | 1.159 | 1.170 | 1.169 |
| BMI | 1.104 | 1.101 | 1.099 | 1.099 | 1.105 | 1.101 | 1.100 | 1.098 |
| SystolicBP | 1.102 | 1.063 | 1.076 | 1.076 | 1.102 | 1.061 | 1.074 | 1.074 |
| DiastolicBP | 1.073 | 1.056 | 1.062 | 1.062 | 1.074 | 1.058 | 1.061 | 1.062 |

Mean *χ*^2^ test statistics across all SNPs (left columns) and genomic inflation factors *λ*_GC_ (right columns) using four methods: LR, linear regression; PCA, linear regression using 10 principal components as covariates; LMM, BOLT-LMM-inf; and BOLT-LMM. In all cases, both mean *χ*^2^ and *λ*_GC_ exceed 1 because of polygenicity. PCA, LMM, and BOLT-LMM are consistently calibrated, whereas uncorrected linear regression suffers inflation due to population stratification, especially for height.

**Supplementary Table 11.** Control for stratification: Height *p*-values at lactase computed by various methods

|  | LR | PCA | LMM | BOLT-LMM |
| --- | --- | --- | --- | --- |
| $\chi^2$ statistic | 33.09 | 3.66 | 6.56 | 6.07 |
| $p$-value | $9 \times 10^{-9}$ | 0.06 | 0.01 | 0.01 |

We report *χ*^2^ statistics and *p*-values computed at rs2011946, the SNP typed in WGHS with highest LD (*R*^2^ = 0.64 in 1000 Genomes reference samples) to lactase-associated SNP rs4988235. Methods: LR, linear regression; PCA, linear regression using 10 principal components as covariates; LMM, BOLT-LMM-inf; and default BOLT-LMM.

**Supplementary Table 12.** Power of mixed model association vs. linear regression for simulated case-control ascertained traits

(a) $h_g^2 = 0.25$ (for liabilities underlying case-control phenotype)
| Method | Case prevalence |  |  |  |  |
| --- | --- | --- | --- | --- | --- |
|  | 50% | 10% | 5% | 1% | 0.1% |
| Linear regression | 2.828 (0.010) | 3.723 (0.015) | 4.401 (0.015) | 6.206 (0.020) | 9.179 (0.026) |
| BOLT-LMM-inf | 2.862 (0.010) | 3.739 (0.015) | 4.384 (0.015) | 5.965 (0.018) | 8.212 (0.023) |
| BOLT-LMM | 2.887 (0.011) | 3.767 (0.015) | 4.397 (0.014) | 5.859 (0.022) | 7.869 (0.025) |

(b) $h_g^2 = 0.5$ (for liabilities underlying case-control phenotype)
| Method | Case prevalence |  |  |  |  |
| --- | --- | --- | --- | --- | --- |
|  | 50% | 10% | 5% | 1% | 0.1% |
| Linear regression | 4.661 (0.014) | 6.452 (0.021) | 7.756 (0.023) | 11.360 (0.030) | 17.290 (0.043) |
| BOLT-LMM-inf | 4.904 (0.016) | 6.582 (0.022) | 7.662 (0.021) | 10.202 (0.025) | 13.252 (0.033) |
| BOLT-LMM | 5.165 (0.018) | 6.798 (0.026) | 7.826 (0.024) | 9.816 (0.034) | 12.326 (0.037) |

We compare mean *χ*^2^ statistics over causal SNPs for linear regression, BOLT-LMM-inf, and BOLT-LMM analysis of simulated case-control traits with prevalences ranging from 50%–0.1%. We simulated genotypes by generating individuals as mosaics of up to 100 random “ancestors” from the WTCCC2 data set, resampling ancestors every 500 SNPs. We restricted the SNP set to the first 2,500 SNPs on each autosome, for a total of M = 55,000 SNPs (so that M_effective_ ≈ 10,000 independent SNPs). We simulated case-control phenotypes using a liability threshold model in which we first generated continuous phenotypes with (a) 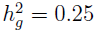 or (b) 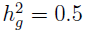 explained by M_causal_ = 1000 markers and then defined cases as individuals with phenotypes exceeding a threshold corresponding to the desired prevalence. In each simulation, we ascertained 5,000 cases and 5,000 controls for a total of N = 10,000 simulated individuals. We then ran association analysis on the ascertained samples. Errors, s.e.m. over 100 simulations.

The results show that for non-ascertained traits, power of linear regression *<* BOLT-LMM-inf *<* BOLT-LMM, consistent with our findings for quantitative traits. This trend reverses for ascertained traits at a prevalence near 5%. For the infinitesimal mixed model vs. linear regression, this result is consistent with the findings of ref. [12], which also observes that the relative reduction in test statistics increases with 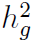 and the ratio N / M_effective_ in addition to ascertainment severity. In our simulations, N ≈ M_effective_, so these simulations correspond to N ≈ 60,000 in real data sets, which have M_effective_ ≈ 60,000 [12]. For data sets with fewer than 60,000 samples, case-control ascertainment will present less of a problem for mixed model methods than indicated above.

**Supplementary Table 13.** Precision of infinitesimal mixed model statistic calibration

| Phenotype | Mean prospective stat | Mean retrospective stat<br>(pre-calibration) | Calibration factor<br>$c_{\text{inf}}$, equation (9) |
| --- | --- | --- | --- |
| ApoB | 1.285 | 1.281 | 0.996 (0.001) |
| LDL | 1.013 | 1.011 | 0.998 (0.000) |
| HDL | 1.026 | 1.024 | 0.997 (0.001) |
| Cholesterol | 0.989 | 0.986 | 0.997 (0.000) |
| Triglycerides | 0.881 | 0.879 | 0.998 (0.000) |
| Height | 0.783 | 0.777 | 0.993 (0.001) |
| BMI | 0.900 | 0.897 | 0.997 (0.001) |
| Systolic BP | 0.853 | 0.849 | 0.996 (0.001) |
| Diastolic BP | 0.770 | 0.769 | 0.998 (0.001) |

We report the BOLT-LMM-inf calibration factor for the nine analyzed WGHS phenotypes. This calibration is similar to GRAMMAR-Gamma [10] but is estimated by computing statistics at only 30 SNPs. The standard errors (jackknife estimates) show that 30 SNPs are enough to achieve high calibration precision.

